# Spatial proteomics reveals lipid droplet reorganization in symbiotic *Paramecium bursaria* cells

**DOI:** 10.1101/2025.05.12.652804

**Authors:** Yan-Jun Chen, Md Mostafa Kamal, Chuan-Chih Hsu, Jun-Yi Leu

## Abstract

Endosymbiosis is an important adaptive mechanism allowing organisms to exploit novel niches. The flexible relationship between the ciliate *Paramecium bursaria* and green algae represents a model system for studying early endosymbiosis evolution. However, the mechanisms underlying how *P. bursaria* cells maintain this endosymbiotic relationship remain unclear. Here, we use mass spectrometry-based proteomics to generate a spatial proteomic atlas of the host *P. bursaria* cells with and without endosymbionts. This atlas defines the protein composition of the endosymbiont-containing compartment (perialgal vacuole) and reveals pronounced remodeling of host lipid droplets upon symbiosis. Imaging analyses confirm that lipid droplets change in size, morphology, and positioning, accumulating near intracellular algae. Perturbing symbiotic cells with chemical inhibitors of lipid metabolism reduces endosymbiotic algal numbers, revealing a role for lipid droplets in the host-endosymbiont interaction. Our data provide a comprehensive resource for protein localization in *P. bursaria* cells and elucidate how those cells remodel an existing cellular compartment to maintain endosymbiosis.

## Introduction

It is widely accepted that an ancient endosymbiotic event between an archaeal cell and an alpha-proteobacterium led to the formation of the eukaryotic cell with mitochondria (*1, 2*). The establishment of endosymbiosis, a close relationship in which one organism (the endosymbiont) lives within another (the host), is often driven by complementation between the endosymbiont and host. Consequently, the interacting partners can explore new niches or resources inaccessible to them when living alone (*3, 4*). Endosymbiosis has played a crucial role during the evolution of eukaryotic cells. In addition to the formation of early eukaryotes, a second endosymbiosis event generated the photosynthetic eukaryotic cell, leading to the evolution of algae and plants (*5, 6*). Since those early events, endosymbiosis has occurred continuously and influenced the evolution of various eukaryotic lineages (*7*). However, little is known about how endosymbiosis is established initially and maintained.

The rapid increase in genomic data in recent years has facilitated the deciphering of pathways potentially involved in host-endosymbiont interactions (*8, 9*). Nonetheless, only a few of them have been validated experimentally. Ciliates have long been known to harbor various endosymbionts, and it has been speculated that such associations enable ciliate populations to dominate in certain marine and freshwater habitats (*10*). Among ciliates, *Paramecium bursaria* is well known for its endosymbiotic relationship with green algae, which has fascinated microbiologists for over a century (*11*). In the wild, most *P. bursaria* cells harbor several hundred algal cells in their cytoplasm, and the endosymbionts can be stably transmitted to daughter cells during cell division (*12*). However, symbiotic *P. bursaria* cells (symbiotic cells) can be artificially induced to lose their algal cells and form aposymbiotic cells (aposymbiotic cells) (*13*). When aposymbiotic *P. bursaria* cells are mixed with free-living *Chlorella* cells, they can quickly reestablish endosymbiosis (*14*). Such flexibility allows scientists to observe and manipulate this endosymbiotic relationship.

Although *Paramecium* species have long served as model systems for cell biology and studies of endosymbiosis (*15*), only a small proportion of their protein-coding genes have been functionally characterized (*16–18*). Even by integrating information on protein domains and sequence homology, potential functions can only be assigned to approximately half of the 15000 protein-coding genes in the *P. bursaria* genome (*19*). Thus, a systematic approach is required to understand the sheer complexity of these ciliate cells.

The emergence of novel subcellular compartments and the subsequent expansion of the associated protein repertoire are key features of eukaryotic evolution. Ciliated protozoa represent a lineage characterized by an abundance of unique subcellular compartments, and they also possess some of the largest cells and gene families among all existing eukaryotes (*20*). In *P. bursaria* cells, many subcellular compartments have been reported as involved in the process of endosymbiosis, with the cytoplasmic space being extensively rearranged to facilitate hundreds of endosymbiotic algal cells (*21*).

Many biological processes are executed in specific organelles or subcellular compartments. Controlling protein abundance and localization enables a cell to regulate these processes accurately. Moreover, cells can remodel or reorganize their subcellular compartments to fine-tune their physiology in response to developmental or environmental cues. Dissecting protein abundance and localization under different conditions can facilitate our understanding of a protein’s potential function and how cells adjust their physiology.

Recent advances in mass spectrometry-based proteomics have substantially improved our understanding of the protein landscape in various cell types (*22, 23*). To gain insights into how *P. bursaria* cells reorganize their subcellular structure to facilitate hundreds of endosymbiotic algae, we generated spatial proteomics maps of aposymbiotic and symbiotic *P. bursaria* cells. Our data reveal a significant increase in the proteins localizing to lipid droplets in symbiotic cells.

Microscopic analysis further showed that the morphology and localization of lipid droplets change close to the endosymbiotic algae. Inhibiting lipid metabolism reduced the number of endosymbionts. Our study not only offers a comprehensive resource of protein localization and translocation data between two types of ciliate cells, but also reveals the involvement of lipid droplets in endosymbiosis.

## Results

### Whole-cell proteomic analysis of *Paramecium bursaria* using differential centrifugation

Although the cell biology of *P. bursaria* has been studied for decades, its genome was only decoded recently (*19, 24*). Nonetheless, a considerable proportion of its encoded proteins have unknown functions. We investigated the spatial distribution of the *P. bursaria* proteome, given that a protein’s subcellular localization can provide information about its function (*25*). Moreover, many subcellular compartments are involved in the process of endosymbiosis (*26–28*). Thus, by comparing proteome maps of aposymbiotic and symbiotic cells, we anticipated gaining insights into how those subcellular compartments are reorganized to facilitate endosymbiosis.

We combined cell fractionation, shotgun proteomics, and machine-learning-based prediction to establish organellar protein localization (*29*). In brief, *P. bursaria* cells were lysed by nitrogen cavitation, and the subcellular compartments were separated by differential centrifugation. We collected 11 fractions, and the distinct protein patterns observed in the individual fractions indicated that our protocol successfully separated different subcellular compartments (fig. S1A and 1B). The collected fractions were subsequently analyzed by liquid chromatography-mass spectrometry (LC-MS) in data-independent acquisition (DIA) mode, and the resulting data were processed with Spectronaut for label-free quantitation (LFQ) analysis (Fig. 1A and data file S1A,B, see Methods for details). We analyzed three biological replicates for both symbiotic and aposymbiotic cells. Protein identification was highly reproducible, with approximately 95% of proteins shared across three replicates in both symbiotic and aposymbiotic cells. Moreover, most proteins presented a low coefficient of variation (CV) between replicates, indicative of high data quality (Fig. 1B and fig. S1C).

**Fig. 1.**
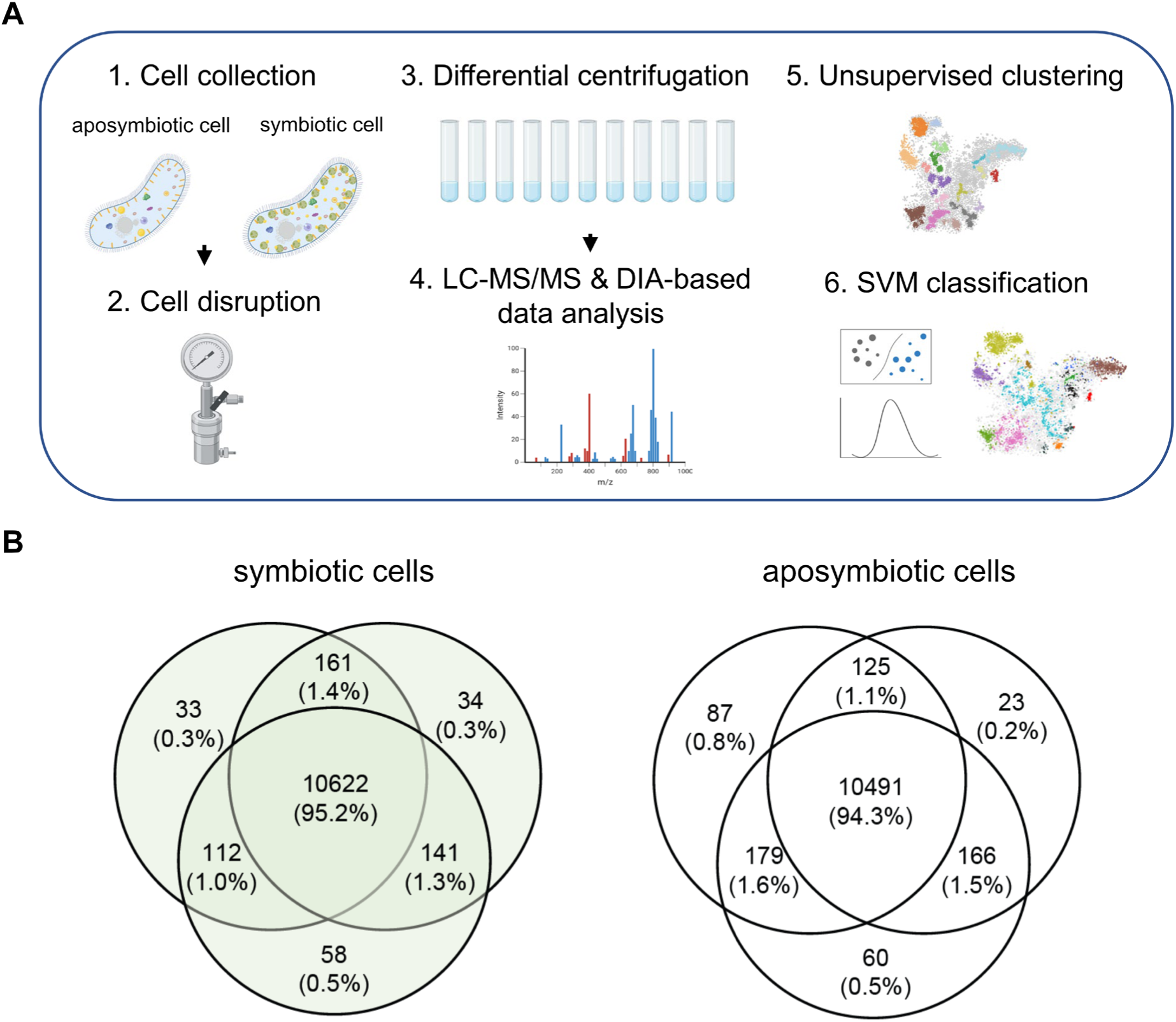
Application of differential centrifugation to separate subcellular compartments of *P. bursaria* cells. **(A)** Workflow of *P. bursaria* subcellular localization mapping by differential centrifugation. *P. bursaria* cells with (symbiotic cell) or without (aposymbiotic cell) endosymbionts were lysed by nitrogen cavitation and the subcellular compartments were separated by differential centrifugation. Subcellular compartment fractions were collected and analyzed by DIA-LC/MS. Protein profiles were generated, subjected to unsupervised clustering, and used to predict protein localizations using SVM machine learning. **(B)** Venn diagrams showing the number of proteins identified in three replicate experiments. The final datasets contained 10622 proteins for symbiotic cells and 10491 proteins for aposymbiotic cells.

In addition to the fractionation samples, we also used mass spectrometry to analyze total cell lysates, allowing us to compare the relative protein abundance of each protein between symbiotic and aposymbiotic cells (data file S2). We detected 6,422 proteins in symbiotic cells and 6,216 proteins in aposymbiotic cells, with 6,135 overlapping between the two cell types. Since our cell lysis protocol was optimized for *P. bursaria* host cells, the endosymbiotic algal cells were not efficiently disrupted. Therefore, the algal proteome is not discussed in this study.

#### Unsupervised clustering maps uncharacterized proteins to subcellular compartments

To globally assess the separation efficiency of different cellular compartments, the fractionation datasets were subjected to unsupervised clustering using hierarchical density-based spatial clustering of applications with noise (HDBSCAN) (*30*). The algorithm evaluated the similarity of protein abundance profiles across 33-plex fractions (from 33 fractions of three replicates) to generate clusters. In total, 5389 proteins in symbiotic cells and 5533 proteins in aposymbiotic cells showed distinct distribution patterns, generating 20 and 19 clusters, respectively (Fig. 2 and data file S3A,B).

**Fig. 2.**
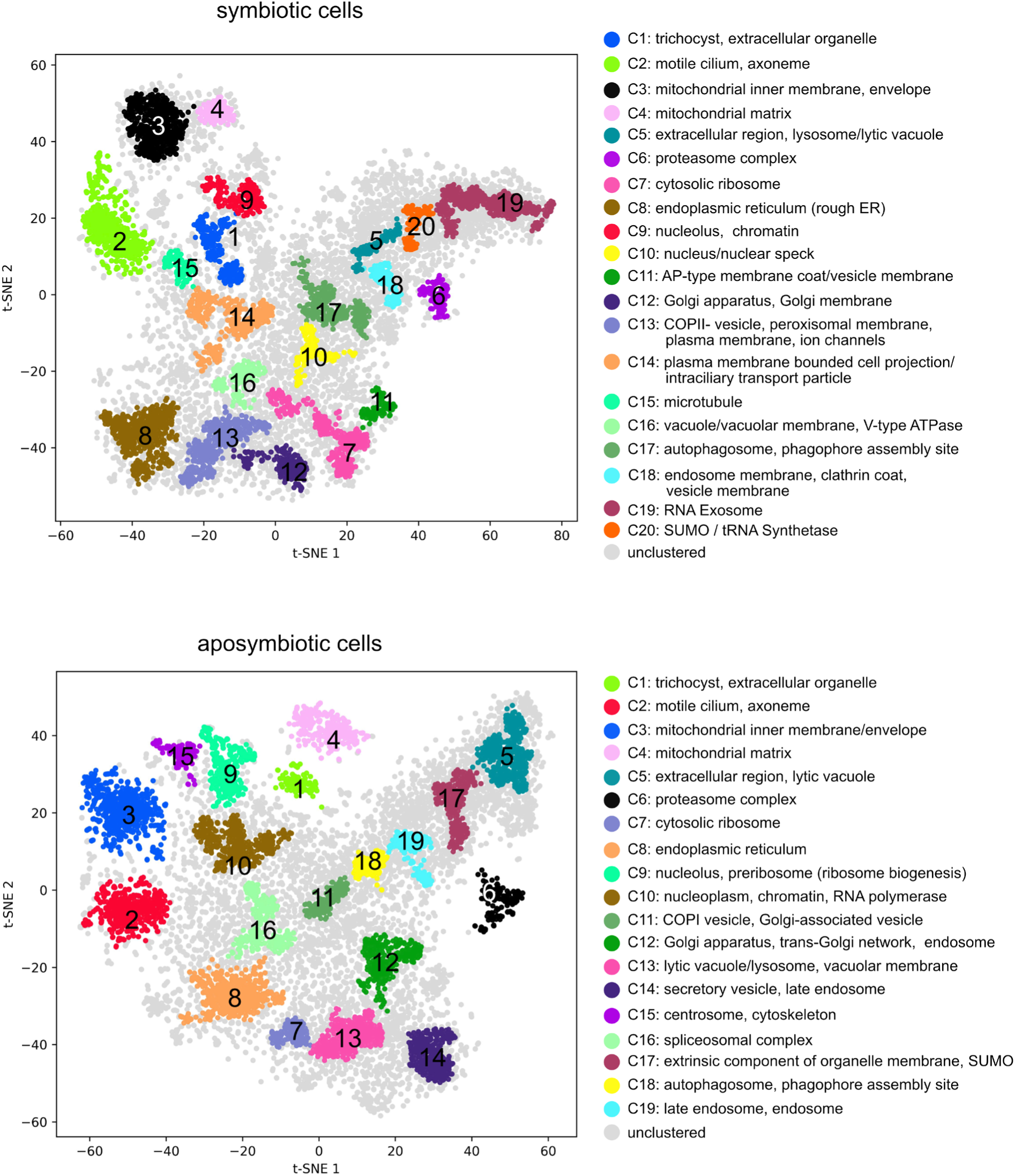
Unsupervised clusters demonstrate a resolution of distinct subcellular functional compartments in *P. bursaria*. HDBSCAN clusters are highlighted on t-SNE projections for **(A)** symbiotic *P. bursaria* cells and **(B)** aposymbiotic *P. bursaria* cells. Each point represents a protein. Most significantly over-represented GOCC terms in each cluster are shown in the right panel, indicating that most clusters have distinct biological functions. In our enrichment test, only terms with an enrichment score greater than 2 and an adjusted *P* value less than 0.05 were selected (see also data file S3C,D).

We utilized several well-conserved marker proteins of proteosomes, ribosomes, and ER, as well as ciliate-specific trichocyst proteins, to examine whether these clusters represent meaningful subcellular niches. These proteins are clearly grouped into clusters 6, 7, 8, and 1, indicating that these subcellular compartments were well separated (fig. S2A). Gene ontology cellular component (GOCC) enrichment analysis further supported that the clusters represent a diverse set of relevant biological niches within the cell (Fig. 2 and data file S3C,D). Clusters 1-12 are highly enriched with GOCC terms related to trychocyst, basal body, mitochondria, lytic vacuole, proteasome, ribosome, ER, nucleus, and Golgi. In some cases, sub-compartment resolution was observed (data file S3C,D). For example, proteins related to the mitochondrial envelope and matrix (clusters 3 and 4 in Fig. 2), as well as those related to the nucleolus and nucleoplasm (clusters 9 and 10 in Fig. 2), were separated into distinct clusters in both cell types. These data indicate that our fractionation data can be used to generate a high-resolution spatial proteomic map. On the other hand, the disruption of cellular compartments during fractionation may lead to divergent distribution patterns among proteins from the same compartment.

Symbiotic cells form a specific membrane compartment, the perialgal vacuole (PV), to accommodate endosymbiotic algae (*31, 32*). We investigated whether one of the HDBSCAN clusters was formed by the proteins of PVs. Several criteria were applied in selecting the candidate PV cluster. First, we focused only on symbiotic-specific clusters whose proteins are not clustered in aposymbiotic cells. Second, we expected that most PV proteins would have higher abundance in symbiotic cells, as aposymbiotic cells do not possess this structure. Third, we expected that the PV cluster should contain a significant proportion of membrane-related proteins. By analyzing all clusters between these two cell types, we found that cluster 18 in symbiotic cells represents a unique cluster, in which proteins were not clustered or were partially clustered in several different groups in aposymbiotic cells (Fig. 3A). Cluster 18 contains 109 protein groups in symbiotic cells. All 109 protein groups are also detected in the aposymbiotic proteome; however, they are dispersed across multiple different clusters rather than forming a single coherent group (Fig. 3A), consistent with the notion that PV proteins are pre-existing host proteins that are relocalized upon symbiosis establishment. We examined whether other clusters also showed differential patterns between symbiotic and aposymbiotic cells. While some clusters exhibited minor differences in protein membership, cluster 18 was the only one that simultaneously met all three of our selection criteria, making it the strongest PV candidate. Moreover, their protein abundance in symbiotic cells is significantly higher than that in aposymbiotic cells (Fig. 3B, C).

**Fig. 3.**
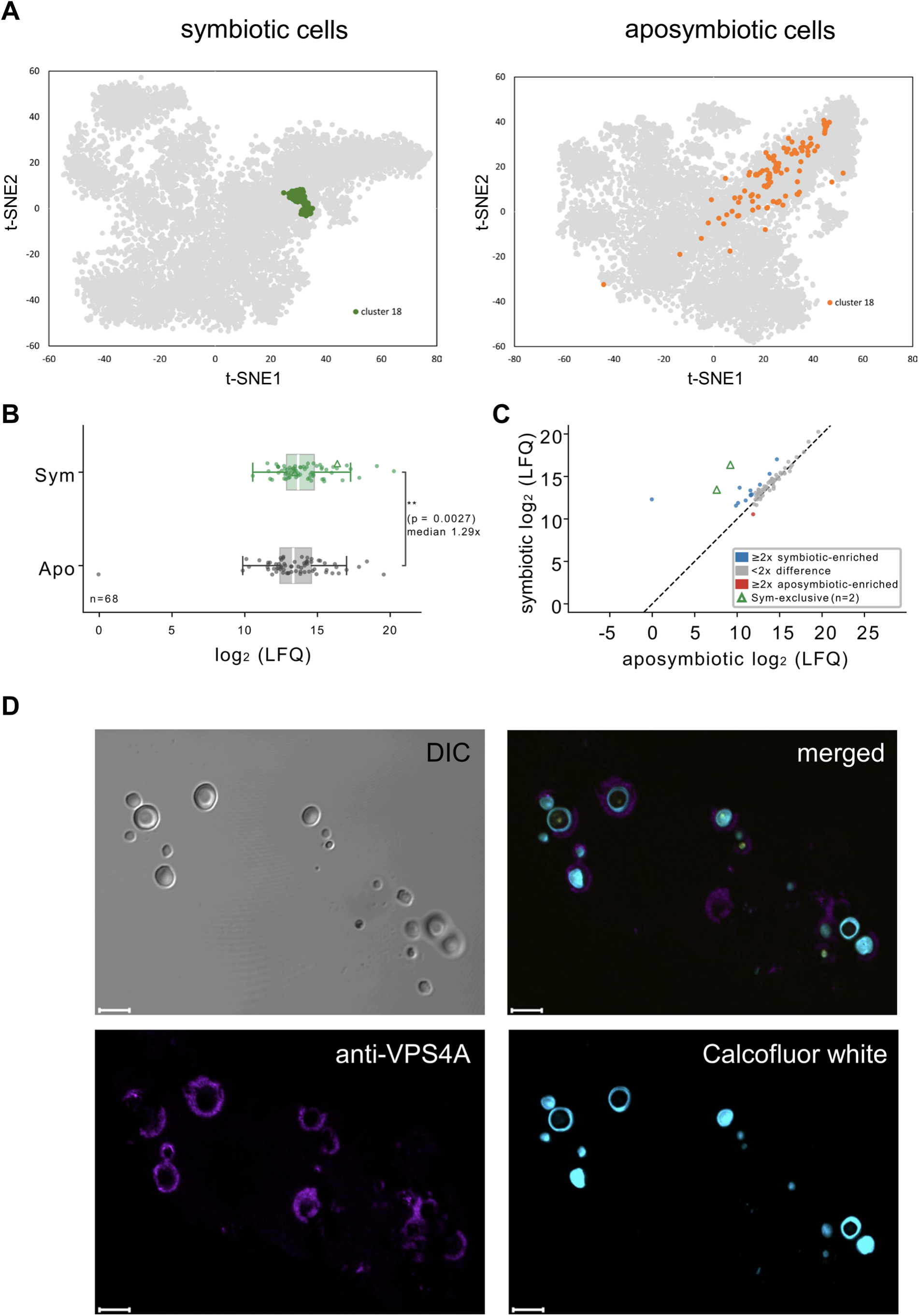
Cluster 18 of the HDBSCAN clusters in symbiotic cells is probably formed by the proteins of perialgal vacuoles. **(A)** The cluster 18 proteins in symbiotic cells are not clustered in aposymbiotic cells. **(B)** The proteins of cluster 18 in symbiotic cells (Sym) have higher abundance than those in aposymbiotic cells (Apo). Protein abundance data were obtained from total lysate proteomes (data file S2). **, *P* value < 0.01, Paired-Sample t-test, n = 68 protein groups. Median fold change, 1.29×. **(C)** Per-protein scatter plot showing the abundance of each cluster 18 protein in symbiotic versus aposymbiotic cells. **(D)** Confocal microscopic images show that VPS4A surrounds the symbiotic algal cell wall. Images represent ultrathin (∼70 nm) resin sections, providing a cross-sectional view of the perialgal region. Calcofluor White stains the algal cell wall as a spatial reference for perialgal localization. See fig. S3 for whole-cell views comparing VPS4A localization in symbiotic and aposymbiotic cells, including specificity controls with food algae. Scale bar, 5 μm.

The proteins in cluster 18 were enriched for GOCC terms including cytoplasmic vesicle membrane, clathrin coat, and endosome membrane (Fig. 2A and data file S3C), which is also consistent with the characteristics of PVs (*27*). Together, these data suggest that cluster 18 is a strong perialgal vacuole candidate.

To experimentally validate the cellular localization of proteins in cluster 18, we performed immunostaining of symbiotic cells with an antibody against one of the cluster 18 proteins, vacuolar protein sorting 4A (VPS4A). The result showed that VPS4A surrounds the algal cell wall inside the symbiotic cell (Fig. 3D), consistent with the expected localization of PVs. In contrast, no VPS4A signal above background was detected in aposymbiotic cells (fig. S3), consistent with the dispersal of cluster 18 proteins across multiple subcellular compartments in aposymbiotic cells (Fig. 3A) and the absence of perialgal vacuole membranes on which VPS4A could concentrate into detectable ring structures. Importantly, in symbiotic cells, VPS4A rings were observed exclusively around endosymbiotic *Chlorella* cells and were absent from ingested food algae (*Chlorogonium capillatum*) undergoing digestion in digestive vacuoles (fig. S3, yellow arrows), demonstrating that VPS4A ring formation is specific to the perialgal vacuole membrane and does not occur on digestive vacuole membranes. In the following analyses, we selected several cluster 18-specific proteins as the PV markers for supervised clustering.

#### A Spatial proteome maps of symbiotic and aposymbiotic *P. bursaria*

To construct spatial maps, marker proteins that collectively represent the cell’s compartments were first required. Since *P. bursaria* is less well studied, the initial marker set was developed based on ortholog localization information from previous studies (*20, 33*) and guided by HDBSCAN clustering (fig. S2B and data file S4A, see Methods for details). Consequently, we selected a total of 252 maker proteins that defined 14 cellular compartments, including the cytosol, proteasome, ribosome, peroxisome, endoplasmic reticulum (ER), Golgi, plasma membrane, nucleus, lipid droplet, basal body, mitochondria, phagolysosome, trichocyst, and potential perialgal vacuole. These cell-compartment markers exhibited distinct fractionation profiles (Fig. 4A and figs. S4 and S5), indicating that the different compartments were well separated.

**Fig. 4.**
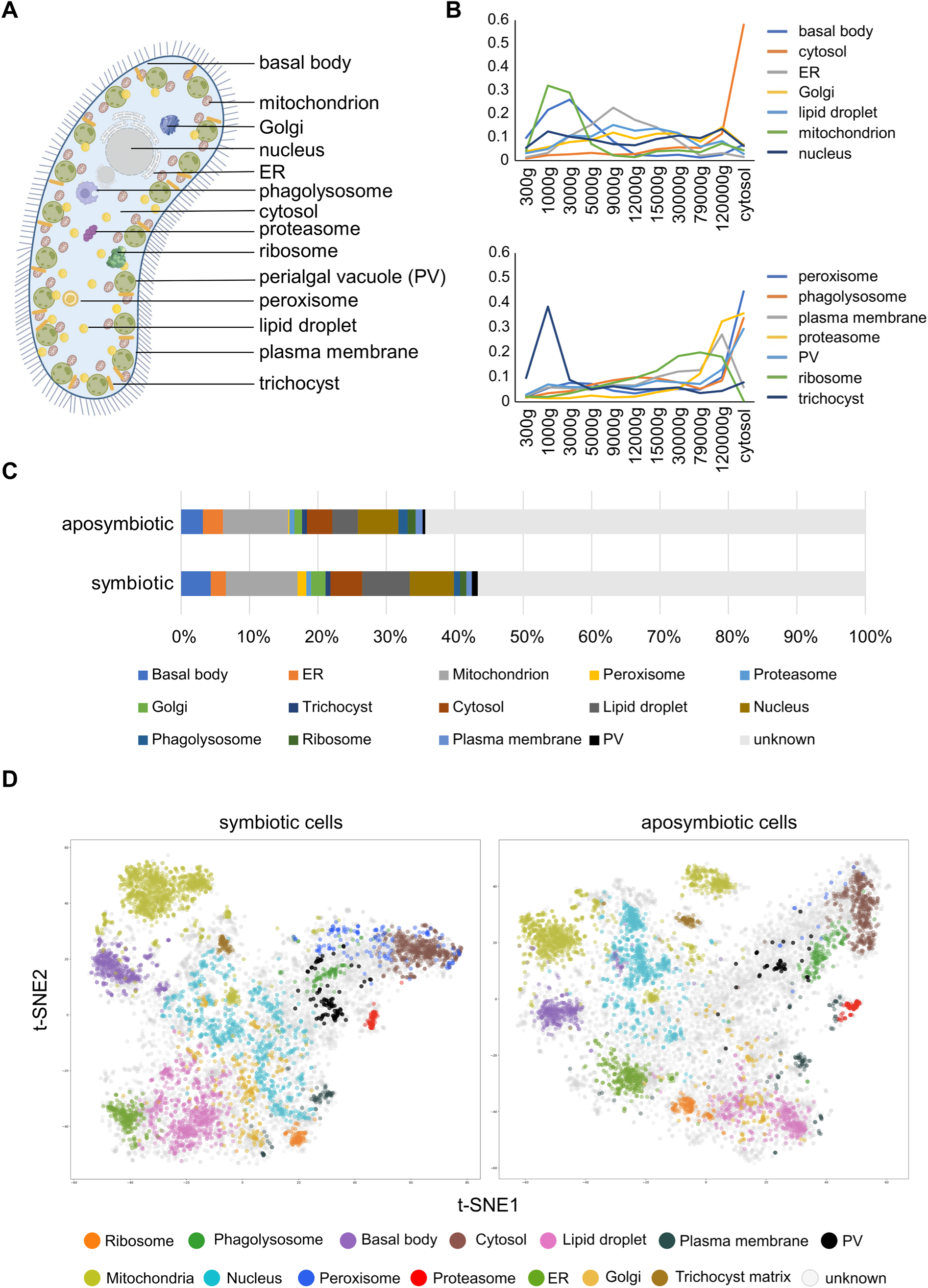
Subcellular localization maps of symbiotic and aposymbiotic *P. bursaria* cells. **(A)** Schematic representation of 14 subcellular compartments in *P. bursaria* cells. **(B)** The distribution profiles of marker proteins. All the marker proteins of the same subcellular compartment were averaged and shown. Distribution profiles of individual marker proteins are shown in fig. S4. Since there were 14 subcellular compartments, we split the distribution profiles into two figures to improve clarity. **(C)** Proportions of identified proteins in different subcellular compartments. Protein localization was assigned using SVM-based machine learning. Only protein assignments with medium or higher confidence are shown. Proteins that could not be assigned to a specific compartment were classified into the “unknown” group. **(D)** t-SNE projections of the proteins assigned to different subcellular compartments in symbiotic *P. bursaria* cells and aposymbiotic *P. bursaria* cells. Each point represents a protein and proteins are colored according to their subcellular localization.

Next, we used the total proteins (n = 10255) that overlapped between aposymbiotic and symbiotic cells for spatial proteome characterization. Our mass spectrometry data were globally normalized and then analyzed using support vector machine (SVM)-based supervised learning, which computes the probability of a protein belonging to one of our pre-defined subcellular compartments by finding the best boundary that separates data points into different classes and maximizing the margin (*34*). Each protein was probabilistically assigned to a compartment.

Allocations represent the most probable compartmental localization of each protein in the dataset. To increase stringency, we retained only protein assignments with medium or higher confidence (i.e., probability > 0.5, see Methods) for further analysis. In total, we classified the subcellular localizations of 4175 and 3387 proteins in the symbiotic and aposymbiotic cells, respectively, representing more than one-third of the identified *P. bursaria* proteome (Fig. 4B, fig. S6, and data file S4B,C). Among the classified proteins, 1237 have not previously had functional annotation. Our data provide additional information on their potential functions.

When we projected the 33-dimension protein profiling dataset (from 33 fractions of three replicates, fig. S7 and data file S1B) of classified proteins into two dimensions by t-distributed stochastic neighbor embedding (t-SNE), we found that the proteins of different cellular compartments formed discrete clusters (Fig. 4C and data file S5A,B), indicating that our analysis can clearly distinguish proteins displaying differential localization patterns. In addition, the distribution of individual subcellular compartments showed highly overlapping patterns with clusters of similar functions in our unsupervised clustering data (Fig. 2 and data file S3A,B).

We further inspected the SVM classification using an alternative localization annotation. Classified proteins from each subcellular localization were subjected to GOCC term enrichment analysis. (Fig. 5A and data file S4D,E, and F). The results revealed consistent and sensible patterns across all cellular compartments in both symbiotic and aposymbiotic cells. For example, trichocyst matrix was enriched with "extracellular membraneless organelle", proteasome with "proteasome complex", basal body with "axoneme" and "motile cilium", and mitochondria with "mitochondrial inner membrane". Notably, although the PV is a symbiosis-specific structure, a small number of proteins with PV-like fractionation profiles were classified into this compartment in aposymbiotic cells (Fig. 5A), likely representing pre-existing endosomal membrane proteins that are reorganized to form the PV upon establishment of endosymbiosis. These analyses demonstrate a high resolution of diverse subcellular compartments with relevant biochemical protein features.

**Fig. 5.**
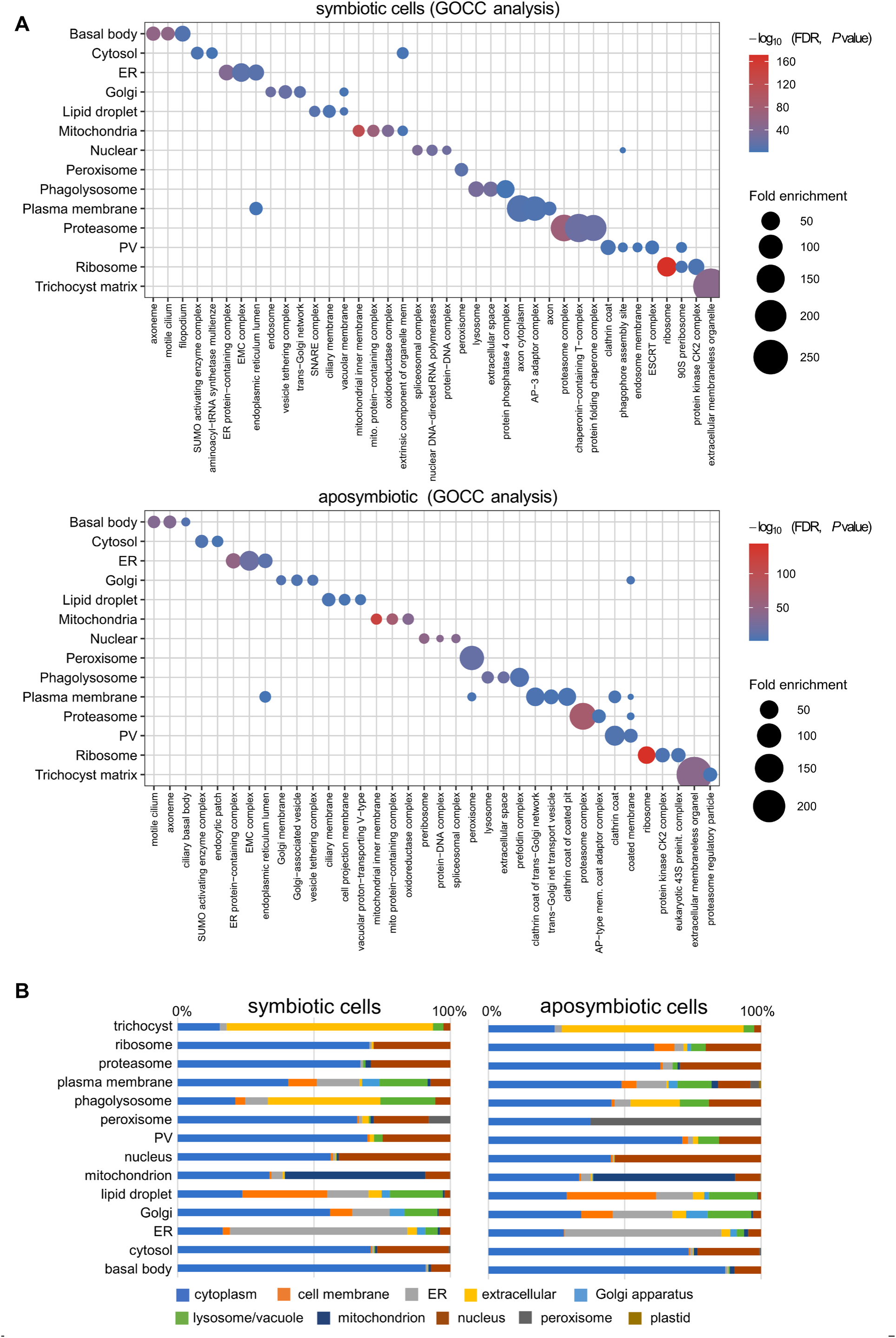
Comparative analysis of spatial proteomes. **(A)** Dot plots of the GOCC term enrichment analysis results for symbiotic (top) and aposymbiotic (bottom) cells. SVM-classified proteins were subjected to GOCC analysis. The Y-axis indicates the subcellular compartments and the X-axis indicates the enriched GOCC terms. Color is scaled by -log₁₀(FDR-adjusted P value) and dot size represents fold enrichment. Aposymbiotic cells lack a true PV; proteins assigned to this compartment by SVM likely represent co-fractionating endocytic membrane proteins. **(B)** Comparison between the results predicted by the pre-trained protein language model, DeepLoc 2.0, and our subcellular localization maps. Color coding represents different predicted subcellular localizations, as determined by DeepLoc 2.0.

#### Comparison of spatial proteomes with deep learning predictions

Since *P. bursaria* cells cannot be genetically manipulated and only a few proteins have been experimentally characterized based on their cellular localization, it is almost impossible to systematically validate our proteomics data. Nonetheless, a pre-trained protein language model, DeepLoc 2.0, has been shown to predict protein subcellular localization with reasonable accuracy (*35*) across several model organisms. We compared our localization data with DeepLoc predictions (data file S6A,B). A large proportion of the proteins assigned to the ER and mitochondria in our experiments were consistently predicted by DeepLoc to localize to those organelles, respectively (Fig. 5B). The majority of nuclear proteins predicted by DeepLoc were also found in a group mapping to the nucleus. Since the training organisms are evolutionarily distant from *P. bursaria*, the accuracy of DeepLoc predictions for ciliate-specific proteins may be reduced. Moreover, approximately one-third of the *P. bursaria* proteome lacks reasonable orthologs in databases of other model organisms.

Interestingly, Deeploc predicted many trichocyst- and phagolysosome-associated proteins as being extracellular. Trichocysts are secretory organelles found in ciliates and some flagellate protists (*36*). They comprise a cavity and long threads of trichocyst proteins that can be released extracellularly in response to external stimuli, such as the detection of predators (*37*).

Phagolysosomes are formed by the fusion of phagosomes with lysosomes after phagocytosis. Although phagolysosomes are primarily involved in degrading food and debris, they may also be used to degrade secreted proteins or exosomes engulfed by phagocytosis or endocytosis (*38*). Further studies of the “extracellular” nature of trichocyst and phagolysosome proteins are needed to understand this finding.

In addition to DeepLoc, we applied a deep-learning protein language model-based algorithm, DeepTMHMM, to predict potential transmembrane proteins (*39*). The analysis showed that proteins assigned to the plasma membrane, lipid droplet, Golgi apparatus, and endoplasmic reticulum (ER) contained a higher proportion of transmembrane proteins (fig. S8 and data file S6A,B), consistent with their membrane-bound or membrane-trafficking roles. In contrast, proteins assigned to the perialgal vacuole showed a comparatively low proportion of predicted transmembrane proteins. However, we note that the high proportion of predicted transmembrane proteins among lipid droplet-assigned proteins likely reflects co-fractionation of ER-resident proteins with lipid droplets, as these two compartments are physically connected through membrane bridges (*40, 41*). Lipid droplets are bounded by a phospholipid monolayer rather than a bilayer (*42*), and many LD-associated proteins use hairpin or monotopic membrane anchors that topology predictors cannot distinguish from true transmembrane helices (*43*). Indeed, recent benchmarking of spatial proteomics methods has shown that lipid droplets are among the most challenging compartments to resolve in density gradient-based approaches, with standard classifiers achieving near-zero performance for this organelle (*44*). Therefore, the SVM-assigned lipid droplet proteome should be considered a broad candidate set that includes both genuine LD proteins and ER-derived co-fractionating proteins. Together, the DeepLoc and transmembrane protein predictions are consistent with the SVM-based classification results. In addition to helping decipher the possible functions of unknown proteins, our comprehensive localization maps enabled us to investigate the physiological changes that occur during the establishment of endosymbiosis.

#### Several cellular compartments are reorganized in symbiotic cells

Given that symbiotic *P. bursaria* cells reorganize their cytoplasm to accommodate hundreds of endosymbiotic algae, we expected that the localization of certain proteins might be altered to accommodate them. Comparing the spatial proteome maps between symbiotic and aposymbiotic cells (Fig. 6A), we observed a dramatic change in protein numbers in the perialgal vacuole (PV), lipid droplet, Golgi, and peroxisome, suggesting that these organelles were potentially reorganized during endosymbiosis. To investigate this observation, we performed Gene Ontology Biological Process (GOBP) enrichment analysis of symbiotic cell-specific proteins in these organelles (data file S7A).

**Fig. 6.**
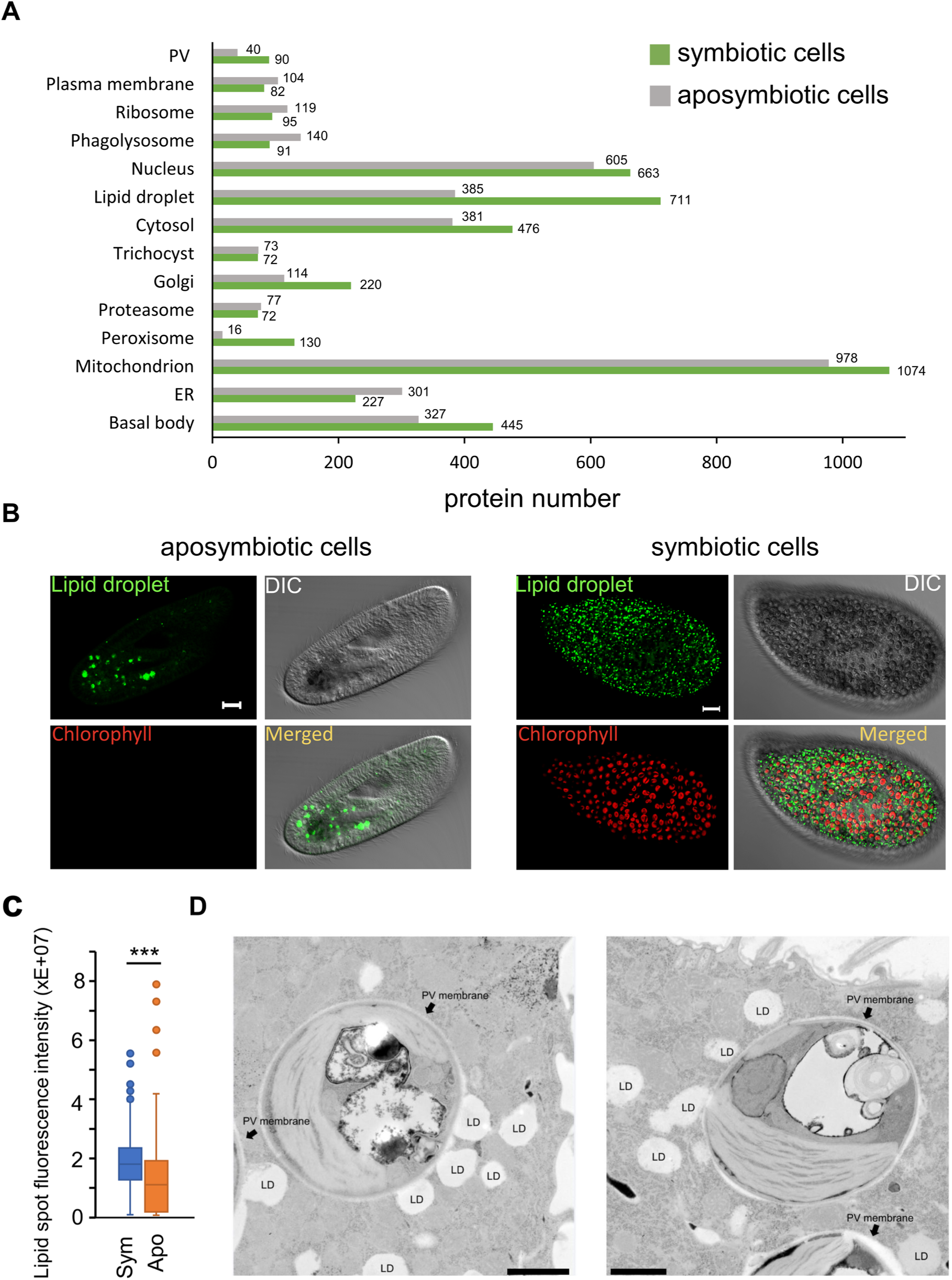
The morphology and localization of lipid droplets are altered in symbiotic cells. **(A)** Numbers of identified proteins that were assigned to different subcellular compartments in symbiotic and aposymbiotic cells. Total protein numbers are indicated next to each bar. **(B)** Lipid droplets in aposymbiotic and symbiotic *P. bursaria* cells. Lipid droplets were stained using LipidSpot (upper panel). **(C)** The LipidSpot fluorescence signal of lipid droplets in symbiotic cells is significantly higher than that of aposymbiotic cells (lower panel). The Cy5 channel was used to detect the chlorophyll of endosymbiotic algal cells. Scale bar, 5 μm. ***, *P* value < 0.001, Mann-Whitney U test, n ≥ 186 cells. **(D)** Many lipid droplets are attached to the perialgal vacuole (PV) membrane of the endosymbionts, as revealed by TEM imaging. The arrow indicates the PV membrane surrounding individual algal cells. Scale bar, 1 μm; LD, lipid droplet.

The perialgal vacuole is a symbiotic cell-specific compartment that protects algal cells from lysosomal degradation in *P. bursaria* cells (*45*). In the symbiotic spatial map, we observed that the PV cluster is located near the phagolysosome cluster (Fig. 4D). Several lysosome-related biological processes are also enriched among symbiotic cell-specific PV proteins (fig. S9 and data file S7A). The enrichment of lysosomal functions in PV proteins is consistent with the hypothesis that PVs may regulate symbiont load through selective digestion of excess endosymbionts.

Notably, a glycogen catabolic process term was also enriched among PV proteins, suggesting potential carbohydrate exchange between the host and endosymbionts at the PV interface. Moreover, many processes related to transport and organelle organization are enriched, consistent with the expected functions of PVs in exchanging nutrients and wastes between the host and endosymbionts.

Lipid droplets regulate intracellular lipid storage, transport, and metabolism (*46*). They are highly dynamic structures, interacting with various organelles (including ER, Golgi, mitochondria, lysosomes, and peroxisomes) to exchange proteins and lipids, and acting as vital hubs of cellular metabolism (*47*). An enrichment analysis of lipid droplet proteins specifically in symbiotic cells revealed enrichment for many pathways involved in lipid metabolism, organelle organization, vesicle trafficking, and ion transport (fig. S9). These pathways are all related to various functions of lipid droplets, indicating that the functions of lipid droplets may shift in symbiotic cells and be involved in the host-symbiont interaction. The enrichment of lipid phosphorylation and lipid transport processes in symbiotic cell-specific lipid droplet proteins suggests active lipid remodeling, possibly to support the metabolic demands of maintaining hundreds of endosymbiotic algae.

Recent studies have highlighted the pivotal role of Golgi-associated mechanisms in regulating symbiotic relationships, as some proteins targeting to the endosymbiont need to be mediated by the Golgi (*48, 49*). In our enrichment analysis of Golgi proteins, we observed that terms related to vesicle trafficking, protein transport, organelle organization, and cellular localization were enriched (fig. S9). The similarity between the enriched terms in symbiotic cell-specific PV, lipid droplet, and Golgi proteins suggests that these cellular compartments may be reorganized to form a central architecture that supports the host-symbiont interaction.

Peroxisomes, traditionally known for their roles in lipid metabolism and reactive oxygen species (ROS) detoxification, are now recognized to respond to pathogens and regulate immune pathways (*50, 51*). However, some viruses and bacteria have evolved mechanisms to hijack this plasticity, remodeling the metabolic output to promote their proliferation (*52, 53*). We observed enrichment in lipid metabolism, stress response, and signaling pathways (fig. S9), which represent the general functions of peroxisomes. The enrichment of oxidative stress response pathways in symbiotic cell-specific peroxisomal proteins is particularly noteworthy, as photosynthetic activity of the endosymbiotic algae is expected to generate elevated levels of reactive oxygen species in the host cytoplasm, necessitating enhanced antioxidant capacity. More experiments are needed to understand the contribution of peroxisomes to symbiotic *P. bursaria* cells.

Previous studies have suggested that mitochondria are involved in the endosymbiosis of ciliates (*26, 54*). It prompts us to examine the mitochondrial proteins more carefully. Interestingly, GO terms related to nucleotide metabolism and stress response were enriched in symbiotic cell-specific mitochondrial proteins, suggesting that mitochondria were under stress (data file S7A). A previous microscopic study showed that symbiotic *P. bursaria* cells contained fewer mitochondria than aposymbiotic ones (*26*). Surprisingly, our total lysate proteomes revealed that the abundance of mitochondrial proteins in symbiotic cells was significantly higher than that in aposymbiotic cells (fig. S10). Since the strain backgrounds are different between these two experiments, it remains to be determined whether the discrepancy is due to the difference in genetic background or methodology.

#### Inhibition of lipid metabolism reduces the number of endosymbionts

Our spatial proteomics data, combined with enrichment analysis, revealed that lipid droplets undergo a dramatic compositional change in symbiotic cells. Consequently, we stained the cells with the lipid droplet dye LipidSpot to examine them by confocal microscopy. We observed that, in individual symbiotic cells, hundreds of tiny lipid droplets seemed to interact with the endosymbiotic algae (Fig. 6B). The total LipidSpot fluorescence intensity of lipid droplets in symbiotic cells was higher than that in aposymbiotic cells (approximately 1.5-fold increase, Fig. 6C). Consistent with this observation, our total lysate proteomics data showed that the protein abundance of several enzymes involved in the biosynthesis of triacylglycerol (TAG), a major core component of lipid droplets, was increased in symbiotic cells (data file S2 and fig. S11).

Next, we used transmission electron microscopy (TEM) to examine the relationship between lipid droplets and endosymbiotic algae in greater detail. Indeed, the lipid droplets were localized near most algae, with some of them (2.4 ± 1.3 lipid droplets per algal cell TEM image, n = 40) being directly attached to the perialgal vacuole (PV) membrane surrounding individual algal cells (Fig. 6D). Together, these observations indicate that the reorganization of lipid droplets in the symbiotic cells may play a role in the host cells’ interactions with their endosymbionts.

We investigated whether lipid droplets are crucial for maintaining endosymbionts using two lipid metabolism inhibitors, H89 and T863. H89 blocks protein kinase A and other kinases, leading to an imbalance between lipolysis and lipogenesis (*55, 56*). T863 can reduce triglyceride synthesis and lipid droplet formation by inhibiting diacylglycerol acyltransferase 1 (DGAT1) (57). We treated symbiotic cells with a mild dose of H89 (1 μM) or T863 (10 μM), which did not affect host survival. Application of these drugs reduced the number of lipid droplets in symbiotic cells (Fig. 7A), and the treatments negatively influenced the number of endosymbionts in host cells (Fig. 7B and 7C). Compared with untreated controls, H89 treatment reduced total algal fluorescence by approximately 40%, whereas T863 treatment reduced it by approximately 30% (Fig. 7C). Together, our data indicate that lipid droplets play a crucial role in endosymbiosis.

**Fig. 7.**
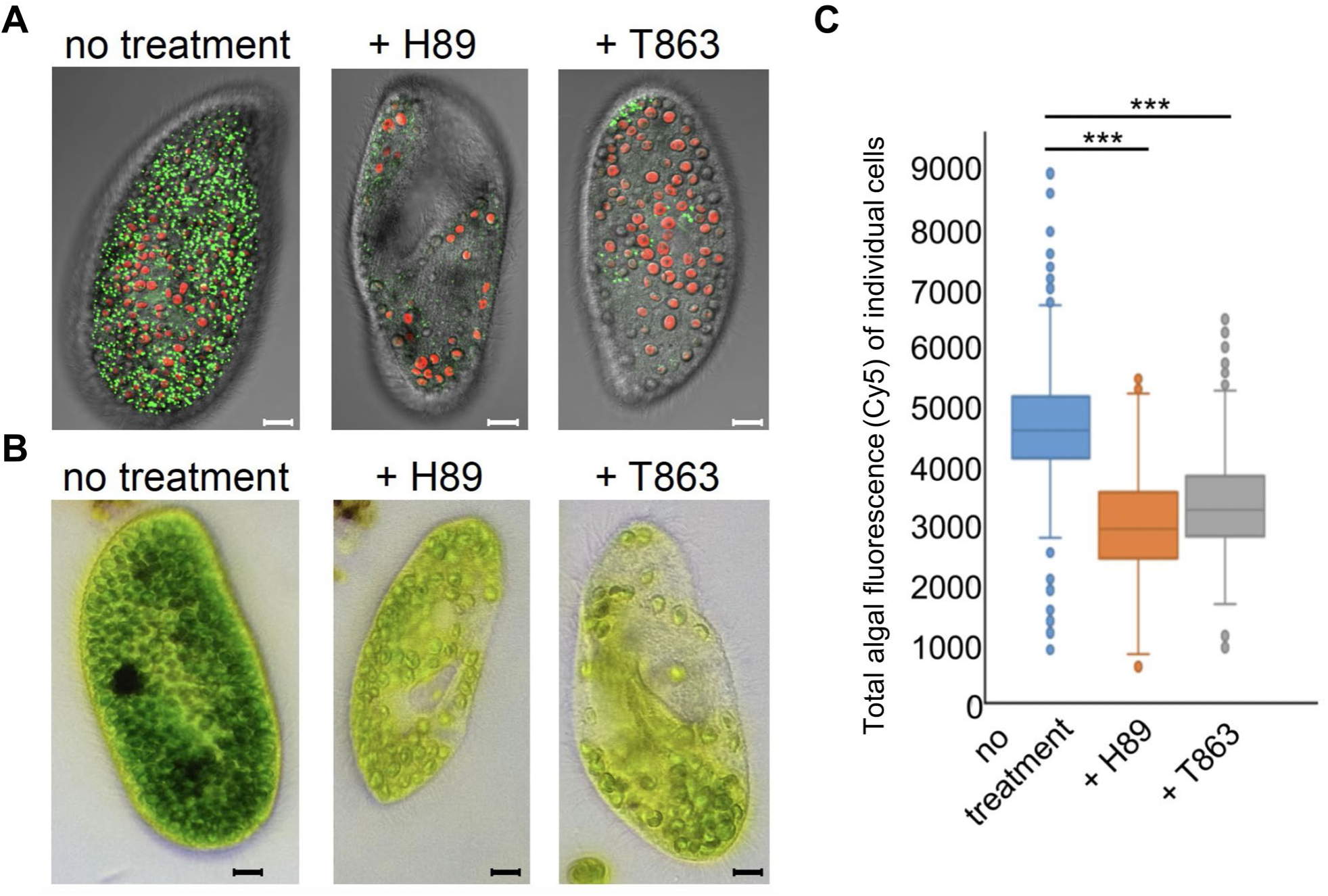
Inhibition of lipid metabolism in symbiotic *P. bursaria* cells reduces the endosymbiont number. **(A)** Treating symbiotic *P. bursaria* cells with mild doses of lipid metabolism inhibitors H89 (1 μM) or T863 (10 μM) reduces the number of lipid droplets. Lipid droplets (in green) were stained using LipidSpot. **(B)** H89 or T863 treatment results in a reduced number of endosymbiotic algae in *P. bursaria* cells. Scale bar, 10 μm. **(C)** Total algal fluorescence (Cy5) of individual cells is significantly reduced after two weeks of treatment. ***, *P* value < 0.001, Mann-Whitney U test, n ≥ 373 *P. bursaria* cells.

## Discussion

*P. bursaria* represents a unique position in the evolution of endosymbiosis in unicellular eukaryotes. In this study, the proteome of two types of *P. bursaria* cells (symbiotic and aposymbiotic) was mapped to 14 subcellular compartments. The fidelity of the classification was verified through multiple orthogonal analyses. Moreover, we identified an important symbiotic cell-specific compartment, the perialgal vacuole, and experimentally validated the localization of one of the predicted PV proteins. Further investigation of the PV proteins will open up opportunities for functional characterization of PVs. We note that while other HDBSCAN clusters also showed minor differences between symbiotic and aposymbiotic cells, cluster 18 was the only one that simultaneously satisfied all three selection criteria (symbiotic-specific clustering, higher protein abundance, and membrane protein enrichment). Notably, the GOCC enrichment of cluster 18 proteins also included food vacuole and phagolysosome terms, consistent with the established model that the PV membrane originates from the host digestive vacuole membrane (*27*), implying that many PV proteins are pre-existing phagocytic membrane components retained during endosymbiosis establishment. Additionally, myosin complex proteins were enriched in cluster 18, suggesting active cytoskeletal remodeling at the PV membrane, a feature that parallels the cytoskeletal remodeling function identified in the cnidarian symbiosome membrane proteome (58). The enrichment of lysosomal functions among PV proteins supports the hypothesis that PVs may participate in symbiont load control through selective digestion of excess algae, a mechanism that would help maintain a stable host-endosymbiont ratio.

The low proportion of DeepTMHMM-predicted transmembrane helices among the classified PV proteins is not inconsistent with enrichment of trafficking and membrane-remodeling factors at the PV, because many proteins that function on endosomal and vacuolar membranes are peripheral rather than integral membrane proteins. Rab GTPases, key coordinators of vesicle identity and trafficking, are prenylated membrane-associated proteins that lack transmembrane helices (*59*). Likewise, ESCRT components are peripheral membrane-associated factors, and VPS4A is a soluble AAA-ATPase that is recruited transiently to ESCRT-III assemblies on membranes rather than spanning the membrane (*60, 61*). Accordingly, the absence of canonical transmembrane helices does not exclude functional association with the PV. DeepTMHMM is a useful screening tool, but negative predictions should be interpreted cautiously for highly divergent proteins from non-model organisms and ideally integrated with orthology, domain content, localization, and experimental data.

The subcellular localization datasets are particularly powerful when comparing different physiological conditions. Our results revealed that proteins assigned to lipid droplets were changed dramatically between symbiotic and aposymbiotic cells. Moreover, lipid droplets underwent morphological changes and often attached to PVs in symbiotic cells (Fig. 6). Inhibition of lipid droplet formation resulted in a significant reduction in endosymbiont numbers, with H89 and T863 treatments reducing total algal fluorescence by approximately 40% and 30%, respectively (Fig. 7). These data suggest that lipid droplets play a role in maintaining endosymbionts. Although the exact boundaries of the lipid droplet proteome are difficult to delineate due to the inherent challenges of resolving lipid droplets in density gradient-based spatial proteomics (*44, 62, 63*), the involvement of lipid droplets in endosymbiosis does not rely on the computational assignments alone. Our confocal microscopy data independently demonstrate altered lipid droplet morphology and increased LipidSpot fluorescence in symbiotic cells, while TEM imaging directly reveals physical attachment of lipid droplets to the perialgal vacuole membrane. Moreover, the upregulation of TAG biosynthesis enzymes in total lysate proteomics confirms elevated lipid metabolism in symbiotic cells at the protein abundance level. Crucially, two pharmacological inhibitors targeting distinct steps of lipid metabolism, H89, which disrupts kinase-mediated lipolysis regulation, and T863, which blocks DGAT1-dependent triglyceride synthesis, both independently reduced lipid droplet numbers and endosymbiont load. This convergence of imaging, biochemical, and functional evidence strongly supports a role for lipid droplets in maintaining endosymbiosis.

Lipid droplets have been observed in several different symbiotic systems and are suggested to perform diverse functions. Symbiotic corals convert lipids or low molecular weight carbohydrates released from dinoflagellate endosymbionts to form gastrodermal lipid bodies (*64, 65*). The lipid composition of lipid bodies varies depending on host physiology (*66*). These lipid droplets likely serve as nutrient storage sites that reorganize the nutrients acquired from the endosymbiont. Nonetheless, they may have additional biological functions that remain to be investigated. It was also observed that endosymbiotic algae in *P. bursaria* enhanced carbon fixation efficiency and accumulated more lipid droplets than free-living algae (*67*). It was suggested that these lipid droplets could serve as an energy source for the host. In a symbiotic bacterial system, *Saccharibacteria* induce their host cells to form lipid droplets (*68*). The accumulation of lipid droplets enables host cells to become more tolerant of environmental stress, suggesting a protective function of lipid droplets. If the function of lipid droplets in *P. bursaria* is similar to the abovementioned cases, the reduced endosymbionts observed in lipid metabolism-compromised cells are probably due to an indirect effect caused by imbalanced nutrient metabolism or weakened host physiology.

Lipid droplets were also observed to play a crucial role in host-pathogen interactions. Pathogens, including viruses, bacteria, and eukaryotic parasites, can manipulate host lipid metabolism and exploit lipid droplets as nutrient sources or as platforms for viral replication (*69*). For example, mammalian cells infected by the protozoan *Toxoplasma* display increased numbers of lipid droplets (*70*). These newly formed lipid droplets attach to the parasitophorous vacuole containing *Toxoplasma* cells, allowing the parasites to utilize the nutrients for their own reproduction (*71*). Similarly, lipid droplets in *P. bursaria* cells may facilitate nutrient uptake by the endosymbiotic algae from the host. In another ciliate symbiotic system, the host and endosymbionts have coevolved a mechanism to balance resource allocation (*54*). Whether remodeled lipid droplets in symbiotic *P. bursaria* cells are used to support the algal growth, as observed in the *Toxoplasma* case, remains to be established.

Since the perialgal vacuole is a membranous structure, lipid droplets may provide essential components for generating or maintaining PVs. Inside *P. bursaria*, a PV membrane surrounds each algal cell, and this membranous structure needs to divide and reorganize each time the algal cells reproduce. If PV structures become compromised, it would likely result in unstable endosymbiosis. Lastly, lipid droplets may serve as a signal-exchange hub between the host and endosymbionts, with their disruption meaning impaired communication between the two interacting partners, potentially prompting digestion of the endosymbionts.

This study establishes a molecular framework for understanding how a ciliate host reorganizes its cellular architecture to accommodate endosymbiotic algae; nonetheless, several aspects invite further investigation. *P. bursaria* is not currently amenable to genetic manipulation, so experimental validation relied on immunolocalization and pharmacological approaches. Among the identified PV candidate proteins, VPS4A is the only one directly confirmed by microscopy, and expanding this set is an important priority for future work. The spatial proteomics data represent a static snapshot of two physiological states and do not capture dynamic protein redistribution during symbiosis establishment or algal division cycles. H89 and T863 have broad cellular targets; while concordant results from two mechanistically distinct inhibitors strengthen our conclusions, indirect effects on the endosymbionts cannot be fully excluded. Finally, this study examined a single laboratory strain, and whether these findings generalize to other *P. bursaria* strains or related ciliate-alga partnerships remains to be determined.

The contribution of lipid droplets to maintaining a mutualistic endosymbiosis adds a new dimension to the already complex functions of these organelles. Peroxisomal proteins involved in oxidative stress response are enriched in symbiotic cells, suggesting that peroxisomes help mitigate reactive oxygen species generated by photosynthetic endosymbiont activity. Together, these findings demonstrate that even unicellular organisms can substantially reorganize their cellular compartments in response to physiological demands. The comparative spatial proteome datasets reported here provide a resource for future studies on ciliate biology and the molecular basis of endosymbiosis.

## Materials and Methods Experimental Design

This study generated comparative spatial proteome maps of symbiotic and aposymbiotic *Paramecium bursaria* cells to identify organelles and proteins reorganized during endosymbiosis. Cells were lysed by nitrogen cavitation and subcellular compartments separated by differential centrifugation, followed by LC-MS/MS proteomics. Protein abundance profiles were analyzed by unsupervised clustering and supervised SVM classification against curated marker proteins to assign subcellular localizations. Total lysate proteomics was performed in parallel to compare protein abundances between cell types. The functional role of lipid droplets in endosymbiosis was assessed by confocal microscopy, transmission electron microscopy, and pharmacological perturbation using lipid metabolism inhibitors.

## Strains and cell culture conditions

All the *P. bursaria* cells used in the study belong to the DK2 strain (*19*). *P. bursaria* cells were cultured in 12 mM sodium acetate, 0.5% yeast extract with modified Dryl’s buffer (2 mM sodium citrate, 1 mM NaH2PO4/Na2HPO4 and 1.5 mM CaCl2) and fed with *Chlorogonium capillatum* (National Institute for Environmental Studies, Japan, strain #NIES-3374) cells every five days.

The cell cultures were kept at 23 °C with a light-to-dark cycle of 12 h:12 h. To obtain aposymbiotic *P. bursaria* strains, green symbiotic cells were treated with 100 μg/ml of methyl viologen dichloride hydrate (Paraquat dichloride, Cat #856177, Sigma-Aldrich, MA, USA) for approximately one week in a light-to-dark cycle of 12 h:12 h (*72*). The aposymbiotic cells were maintained under low-light conditions.

For the inhibitor treatment experiments, 20 ml of symbiotic *P. bursaria* cells (1000 cells/ml) were treated with different concentrations of H89 (Cat. # HY-15979, MedChemExpress, NJ, USA) or T863 (Cat. # HY-32219, MedChemExpress) to test their viability and growth. We used 1 μM of H89 and 10 μM of T863 in the final experiment since these drug concentrations did not have any apparent deleterious effects on *P. bursaria* cells. After two weeks of treatments, we observed the morphology of the host lipid droplet and symbiotic algae by confocal microscopy.

## Total lysate preparation and subcellular compartment fractionation

For each biological replicate of the total proteome experiment, 1 × 10⁵ *P. bursaria* cells (approximately 0.5 L of the culture volume) were harvested on an 11-μm-diameter nylon mesh sieve after removing cell debris with cheesecloth. The cells were washed on the nylon mesh with Dryl’s solution and then transferred to 1.5-ml tubes. The cells were resuspended in 1 ml of UTU buffer (6 M urea, 2 M thiourea, 100 mM Tris, 2.5 mM DTT, 5 mM EDTA-KOH [pH 8.0], 1 mM/2 mM PMSF/SHAM, and 1× protease inhibitor cocktail). After centrifugation at 12,000 × g, the supernatant was collected for LC-MS analysis.

For each biological replicate of the subcellular fractionation experiment, 4 x10^5^ *P. bursaria* cells (around 2 L of the culture volume) were harvested from an 11 μm-diameter nylon mesh sieve after removing cell debris using cheesecloth. The cells were washed on the nylon mesh with Dryl’s solution before being decanted into 15 ml tubes. Then, the cells were resuspended in lysis buffer (10 mM HEPES-KOH [pH 7.5], 2 mM EDTA-KOH [pH 8.0], 2 mM magnesium, 1 mM/2 mM PMSF/SHAM, 250 mM sucrose and 1× protease inhibitor cocktail) and lysed using a nitrogen cavitation bomb. The cavitation bomb was pressurized to 300 psi and incubated for 20 min before slowly releasing the pressure.

The lysate was then differentially centrifuged at 300 x *g* for 3 min, 1000 x *g* for 10 min, 3000 x *g* for 10 min, 5000 x *g* for 10 min, 9000 x *g* for 15 min, 12000 x *g* for 15 min, 15000 x *g* for 15 min, 30000 x *g* for 20 min, 79000 x *g* for 43 min, and 120000 x *g* for 45 min. After each centrifugation, the pellet was collected and resuspended in UTU buffer and stored at -80 °C. After the 12000 x *g* centrifugation step, we added acetone to the supernatant to precipitate the protein overnight at -20 °C. Protein concentration was determined by Bradford assay.

Fractions (5ug loading) were analyzed by Western blotting using a standard method with antibodies against the protein marker. The antibody against a *P. bursaria* mitochondrial protein (ubiquinol-cytochrome c reductase complex protein) was generated by inoculating a rabbit with a synthesized peptide. The antibody was diluted to 1μg/ml for immunoblotting. Protein bands were visualized via the chemiluminescence detection using Western Lightning Plus-ECL (PerkinElmer, CT, USA).

## Sample preparation and LC-MS analysis

Protein digestion in the S-Trap microcolumn (ProtiFi, NY, USA) was performed according to a previous report (*73*). In brief, 5 μg of protein in lysis buffer was reduced and alkylated using 10 mM Tris(2-carboxyethyl) phosphine hydrochloride and 40 mM 2-chloroacetamide at 45 °C for 15 min. A final concentration of 5.5% (v/v) phosphoric acid (PA), followed by a six-fold volume of binding buffer (90%, v/v, MeOH in 100 mM triethylammonium bicarbonate (TEAB)), was next added to the protein solution. After gentle vortexing, the solution was loaded into an S-Trap microcolumn. The solution was removed by spinning the column at 4,000 x *g* for 1 min. The column was washed three times with 150 μl binding buffer. Finally, 20 μl of digestion solution containing 200 ng Lys-C and trypsin in 50 mM TEAB was added to the column and incubated at 47 °C for 2 h. Each digested peptide was eluted using 40 μl of three buffers consecutively: (1) 50 mM TEAB; (2) 0.2% (v/v) formic acid (FA) in H2O; and (3) 50% (v/v) acetonitrile (standard elution). Elution solutions were collected in a tube and dried under a vacuum. A total of 500 ng peptides was loaded into an Evotip and then the Evotip was placed into an Evosep One system (Evosep Biosystems, Odense, Denmark) coupled to a timsTOF HT mass spectrometer (Bruker Daltonics, MA, USA).

Indexed retention time peptides (iRT) (KI-3002-2, Biognosys, Schlieren, Switzerland) were dissolved in 50 μl of 0.1% (v/v) FA with 3% (v/v) ACN to achieve a 10× stock solution. The stock solution was then diluted 10-fold. Dried peptides were reconstituted in 200 μl of 0.1% (v/v) trifluoroacetic acid. Then, 20 μl of samples and 10 μl of 1× iRT solution were loaded onto EvoTips and analyzed using the same Evosep One system.

Eluting peptides were separated analytically using the 30 SPD method on an Evosep Performance column (15 cm x 150 µm ID, 1.5 µm, EV1137; Evosep) at 40 °C within a Captive Spray source (Bruker Daltonics), equipped with a 20 µm emitter (ZDV Sprayer 20, Bruker Daltonics), followed by dia-PASEF mode analysis. For dia-PASEF, the instrument firmware was adapted to perform data-independent isolation of multiple precursor windows within a single TIMS frame. An optimized dia-PASEF scheme was applied that targeted +2 and +3 ions in a two-window method. Sixteen of these scans (resulting in 32 windows, each of 25-Da window size) covered an m/z range of 400 to 1200. Each dia-PASEF cycle comprised an MS1 survey frame, followed by 16 dia-PASEF frames, with this setup resulting in a total cycle time of 1.8 s.

## Data-independent acquisition (DIA) analysis

DIA data signal extraction and quantification were performed with an analysis pipeline in Spectronaut (Biognosys, v18.7, https://biognosys.com/software/spectronaut/) using a standard setting with Carbamidomethylation (Cys) specified as a fixed modification, oxidation (Met), and N-terminal acetylations specified as variable modifications. We selected dynamic retention time prediction with local regression calibration. Interference correction for the MS and MS2 levels was enabled. The FDR was set to 1% at peptide precursor and protein levels using scrambled decoy generation and a dynamic library size fraction set at 0.1. MS2-based quantification was used, enabling local cross-run normalization. Peptide precursor abundances were measured by the sum of fragment ion peak areas and peptide grouping was performed based on modified peptides. Protein-level quantification was performed against the spectrum library generated *in silico* by Spectronaut with standard settings and a stripped peptide sequence area obtained from the mean precursor area.

## Data processing and assessment of proteomics depth

For label-free quantitation (LFQ) maps, Spectronaut already globally normalizes intensities, hence no further normalization was required. Proteins not identified in all replicates were removed. Three biological replicates were performed for each cell type. Proteomic differences were evaluated for statistical significance (*P*<0.05) by Student’s t-tests and corrected for multiple testing using the Benjamini-Hochberg correction. Normalization (0-1) of each fractionation profile was performed by summing the intensities across all fractions and dividing each intensity by the profile sum. Proteomics depth was assessed by counting protein groups or proteins identified in one or all replicates.

## Unsupervised clustering

Normalized protein abundance profiles from 33 fractions of three replicates were concatenated and analyzed using hierarchical density-based spatial clustering of applications with noise (HDBSCAN) (*30*). The algorithm generated 20 and 19 clusters in symbiotic and aposymbiotic cells, respectively. The HDBSCAN algorithm is available in Python Scikit-learn (*33*).

## Marker protein curation and support vector machine (SVM) analysis

To generate the subcellular compartment marker protein list, we first examined the marker proteins identified in *Paramecium tetraurelia* (*74*) by manually reviewing the literature and InterProScan annotations. Next, we used OrthoFinder (*75*) to identify orthologous markers in the *P. bursaria* genome. These were then further selected guided by HDBSCAN unsupervised clustering. These marker proteins were inspected for their protein abundance distribution profiles and checked for their deviation from the cluster using a 5-fold cross-validation function provided in Scikit-learn (*33*). Only the marker proteins present in all compared datasets were included, and identical SVM parameters were used.

Classification of unlabeled proteins was performed using SVM. Based on SVM, we achieved a prediction accuracy (F1-score) of 95.2% for symbiotic and 91.5% for aposymbiotic cells (fig. S5a). Finally, we established a list of manually curated proteins defining 14 subcellular compartments (cytosol, proteasome, ribosome, peroxisome, ER, Golgi, plasma membrane, nucleus, lipid droplet, basal body, mitochondrion, phagolysosome, PV membrane and trichocyst) (data file S4A). To further evaluate the performance of our subcellular compartment maps, their power to predict protein localization was assessed using quality-filtered and normalized (0-1) data with full replicate coverage.

For supervised classification, the marker proteins covering 14 subcellular localizations were used by SVM to assign all other proteins to the organellar cluster. Machine learning was conducted using the functionality-based command provided in Scikit-learn (*33*). SVM enables selection of marker classes, definition of a hold-out test set, automated hyperparameter optimization and, finally, SVM training and prediction. During training, the parameters C and gamma of the radial basis function were optimized via an iterative grid search, with 5-fold cross-validation. At each grid point, SVMs were run on each loaded dataset before identifying the optimum for summed accuracy across datasets. Finally, we confirmed analytical performance using Q-Sep (*76*), which quantifies the separation between a pair of predicted protein sets by comparing intra-group and inter-group distances (fig. S5b and data file S8A,B).

For prediction, the SVM was fitted using the training set and 5-fold cross-validation to convert the raw SVM scores into probabilities. Proteins were then assigned to the best-fitting organelle model and divided into confidence classes based on probability: >0.9, very high confidence; >0.8, high confidence; >0.5, medium confidence; >0.4, low confidence; <0.4, best-guess assignment (*77*). In our dataset, probabilities lower than 0.5 were labeled as unknown. For the hold-out test set, a misclassification matrix was derived to calculate global marker prediction recall (proportion of correctly predicted markers to the total number of markers), organelle-specific recall (proportion of markers correctly assigned to the cluster), and organelle-specific precision (ratio of markers correctly assigned to the number of all markers assigned to the cluster). The harmonic mean of recall and precision (F1 score) was used as the primary readout for SVM performance.

## t-SNE (t-distributed stochastic neighbor embedding) visualization

t-SNE was used to reduce dimensionality and visualize data. For t-SNE, the preprocessed data were embedded into two dimensions using a perplexity of 60 and extract gradient calculation for a maximum of 1000 iterations. The computed coordinates were recorded and used to obtain the two-dimensional data projection presented herein.

## Protein annotation and GO/KEGG enrichment analysis

GO terms and KEGG terms were predicted by InterProScan (*78*), the String database 12 (*79*), DeepFRI (*80*), and Pannzer 2 (*81*). Fisher’s exact tests were applied to the annotation terms. The resulting *P* values were corrected for multiple hypotheses using the Bonferroni correction, and a cutoff of 5% FDR was applied (*82*).

## DeepLOC and DeepTMHMM predictions

For subcellular localization, protein feature annotations were predicted using DeepLoc 2.0 (*35*). Transmembrane span prediction was performed using DeepTMHMM (*39*) in *P. bursaria* annotated protein sequences.

## Confocal microscopy

*P. bursaria* cells were subjected to various staining procedures to visualize lipid droplets, cell walls, and protein localization. All samples were prepared under consistent fixation and imaging conditions to ensure comparability across experiments. For lipid droplet staining, cells were similarly fixed with 4% formaldehyde for 10 minutes and stained with LipidSpot dye (1000X dilution; Cat. # 70065-T, Biotium, CA, USA) for 30 minutes at room temperature. Fluorescence signals were detected using a confocal microscope with an excitation wavelength of 440 nm and an emission wavelength of 585 nm.

For immunofluorescence detection of VPS4A, cells were fixed in 4% paraformaldehyde in PBS using rapid microwave irradiation (PELCO 3451 Laboratory Microwave System, Ted Pella) with the ColdSpot set to 10 °C. The program consisted of three cycles (1 min on - 1 min off - 1 min on - 1 min off - 1 min off - 1 min on) at 250 W. After fixation, samples were rinsed with PBS and dehydrated through a graded ethanol series using microwave-assisted processing (one cycle: 1 min on - 1 min off - 1 min on at 100 W). Dehydrated samples were infiltrated with a graded series of HM20 resin (3 minutes per step at 250 W), embedded in gelatin capsules, and polymerized under UV light. Ultrathin ∼70 nm sections were prepared using a Leica EM UC6/UC7 ultramicrotome. Sections were blocked with 1% BSA in PBS for 20 minutes, followed by overnight incubation at 4 °C with a primary anti-VPS4A antibody (Cat# 14272-1-AP, Proteintech, IL, USA). After three PBS washes, sections were incubated with an Alexa Fluor 488-conjugated anti-rabbit secondary antibody (Cat# A-11008, Invitrogen, MA, USA). Imaging was performed using a ZEISS LSM 780 Upright Confocal Microscope (ZEISS, Oberkochen, Germany). Calcofluor White binds to chitin and cellulose present in algal cell walls but does not stain *P. bursaria* host cell structures, providing specificity for identifying the algal cell boundary within the host cytoplasm. For cell wall staining, ultrathin sections were stained with Calcofluor White stain (Cat# 18909, Sigma-Aldrich) for 5 minutes and then rinsed with PBS prior to imaging.

To obtain quantitative measurements of algal autofluorescence (Cy5 channel) and lipid droplet signals at scale, we employed the ImageXpress® Confocal HT.ai High-Content Imaging System (Molecular Devices, CA, USA), this system is equipped with an automated autofocus function, which selects the optically brightest and sharpest focal plane for each field of view—ensuring consistent imaging quality across hundreds of cells, which enabled high-throughput scanning and analysis of hundreds of *P. bursaria* cells under symbiotic and aposymbiotic conditions.

## Transmission electron microscopy (TEM) imaging

*P. bursaria* cells were collected and fixed in a mixed aldehyde solution containing 2.5% glutaraldehyde (GA) and 4% paraformaldehyde (PFA) in 100 mM sodium cacodylate buffer (pH 7.2). Fixation was performed using a microwave. Samples were irradiated at 500W for 1 min, followed by a 30-s pause, and then an additional 1-min irradiation under vacuum (20 in. Hg). This cycle was repeated eight times. After the final microwave cycle, samples were shaken for 30 min at room temperature, followed by an additional 2 h of shaking before overnight storage at 4 °C.

Following fixation, samples were rinsed in 100 mM sodium cacodylate buffer (pH 7.2) under microwave conditions of 150W for 1 min, repeated four times, with no vacuum applied. Post-fixation was performed using 1% osmium tetroxide (OsO₄) in 100 mM sodium cacodylate buffer (pH 7.2). The post-fixation process followed similar microwave conditions: 450W for 1 min, followed by a 30-s pause, and another 1-min irradiation under a 20-in. Hg vacuum. This cycle was repeated eight times, with samples shaken at room temperature for 30 min after each cycle, followed by a final 2-h shaking period before overnight storage at 4 °C.

After post-fixation, samples were rinsed with distilled water (ddH₂O) under microwave conditions of 150W for 1 min, repeated four times, without a vacuum. En-bloc staining was performed using 2% uranyl acetate (UA). The staining process was carried out under microwave irradiation at 450W for 1 min, followed by a 30-s pause, and another 1-min of irradiation with a vacuum of 20 in. Hg. After staining, samples were rinsed with ddH₂O, with four cycles of microwave irradiation at 150W for 1 min per cycle, without a vacuum. Samples were then dehydrated through a graded ethanol series (10%, 20%, 30%, 40%, 50%, 60%, 70%, 80%, 90%, and 100%). Each ethanol step involved microwave irradiation at 200W for 10 s, followed by a pause of 20 s, and another 10-s irradiation. This process was repeated twice for each ethanol concentration, without a vacuum.

Following dehydration, resin infiltration was carried out using spurr’s resin without an accelerator. Initially, 5% spurr’s resin was applied for 2 days at room temperature. Samples were then infiltrated through a graded resin series (10% to 100%) under a vacuum, with shaking every 8 hours. Two changes of 100% spurr’s resin without accelerator were applied at 8-h intervals under a vacuum, followed by three changes of 100% spurr’s resin with the accelerator under the same conditions. Finally, resin-infiltrated samples were embedded in molds and polymerized in a pre-warmed oven at 70 °C for 24 hours.

## Statistical Analysis

Statistical tests used in this study are described in the corresponding Methods subsections and figure legends. Unless otherwise noted, P values from enrichment analyses were corrected for multiple comparisons using the Bonferroni correction with a 5% FDR cutoff. Proteins with a missing value in any fraction were excluded from classification analysis. No statistical methods were used to predetermine the sample size. The experiments were not randomized and the investigators were not blinded to allocation during experiments and outcome assessment.

## Acknowledgments

We thank members of the Leu lab for helpful discussion and comments on the manuscript. We also thank John O’Brien for manuscript editing, the IMB Imaging Core for imaging experiments, and the IPMB Proteomics Core for proteomics experiments. We thank Professor Toshinobu Suzaki (Kobe University, Japan) for helping us set the *Chlorogonium capillatum* feeding axenic method for *P. bursaria* cell culture. We would like to dedicate this paper to the memory of Dr. Meng-Chao Yao (1949-2025), a great mentor and excellent ciliate biologist, who inspires us to work on ciliates

## Author contributions

Conceptualization: MMK, JYL Methodology: YJC, MMK, JYL Investigation: YJC, MMK, CCH Formal analysis: YJC, MMK, CCH Data curation: YJC, MMK Software: YJC, MMK Visualization: YJC, MMK Resources: JYL

Supervision: JYL

Writing-original draft: YJC, MMK, JYL

Writing-review & editing: YJC, MMK, CCH, JYL

**Competing interests:** The authors declare that they have no competing interests.

## Data and materials availability

The mass spectrometry proteomics data have been deposited in the ProteomeXchange Consortium under accession numbers PXD055842 and PXD055875. Additional data generated or analyzed in this study are provided as Data files and Source Data files with this article.

