## supplementary figures for "Spatial proteomics reveals lipid droplet reorganization in symbiotic *Paramecium bursaria* cells"

Yan-Jun Chen *et al.*

**This PDF file includes:**

Figs. S1 to S11

Data S1 to S8

References (# to #) (if applicable—these should refer only to references in the SM)

### Supporting figures

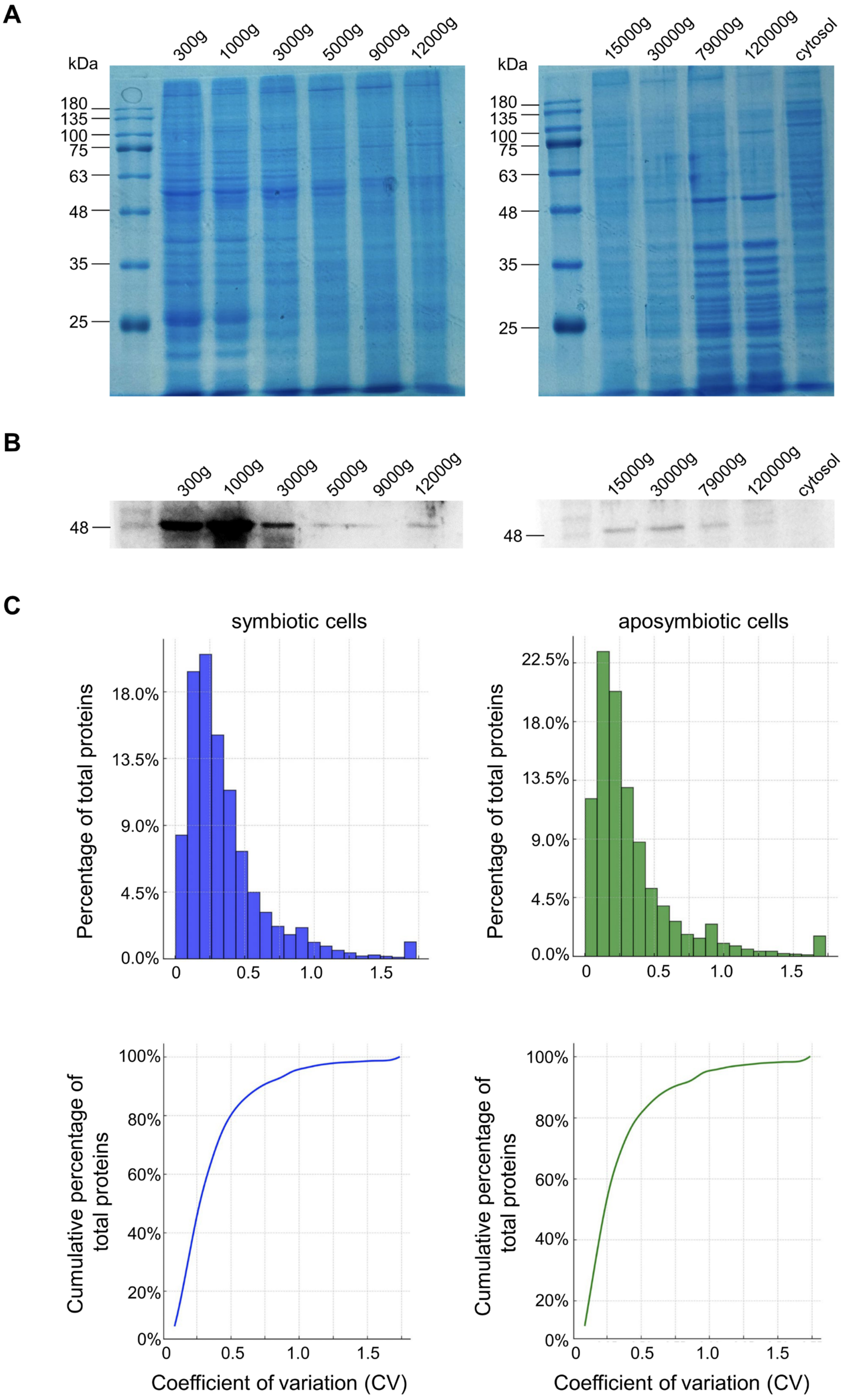

**Fig. S1. Differential centrifugation consistently separates different populations of proteins.** **(A)** SDS-PAGE of subcellular fractions collected from differential centrifugation. **(B)** The Western blot image of anti-ubiquinol-cytochrome c reductase complex protein (a mitochondrial protein, M.W. = 52 kDa) staining in differential centrifugation fractions. **(C)** Proteins identified from three replicates exhibit small coefficients of variation (CV) in terms of their abundance. Only proteins identified in both symbiotic and aposymbiotic cells were analyzed (n = 10255). The top panel shows the distribution of protein abundance CV, and the bottom panel shows the cumulative percentage of total protein abundance CV. The medians of symbiotic and aposymbiotic cells are 0.25 and 0.21, respectively.

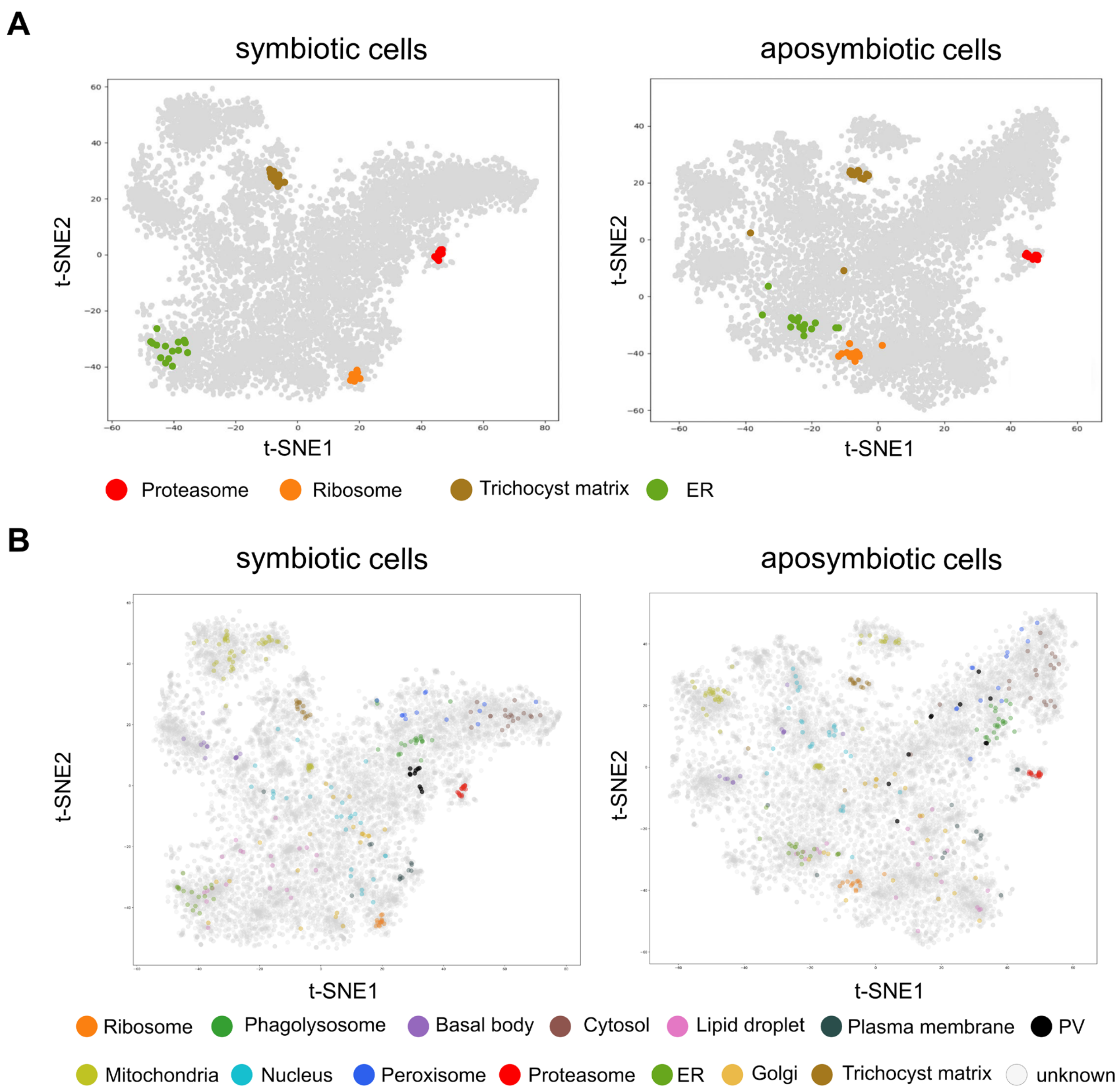


**Fig. S2. Marker proteins of different subcellular compartments are well separated.**
**(A)** Marker proteins in proteosomes, ribosomes, trichocyst, and ER are clearly grouped into the HDBSCAN clusters 6, 7, 8, and 1. **(B)** t-SNE projections of all marker proteins. The markers of some organelles (e.g., the mitochondrion and nucleus) were separated into different clusters, probably due to the breaking of organelles, as indicated by unsupervised clustering (Fig. 2).

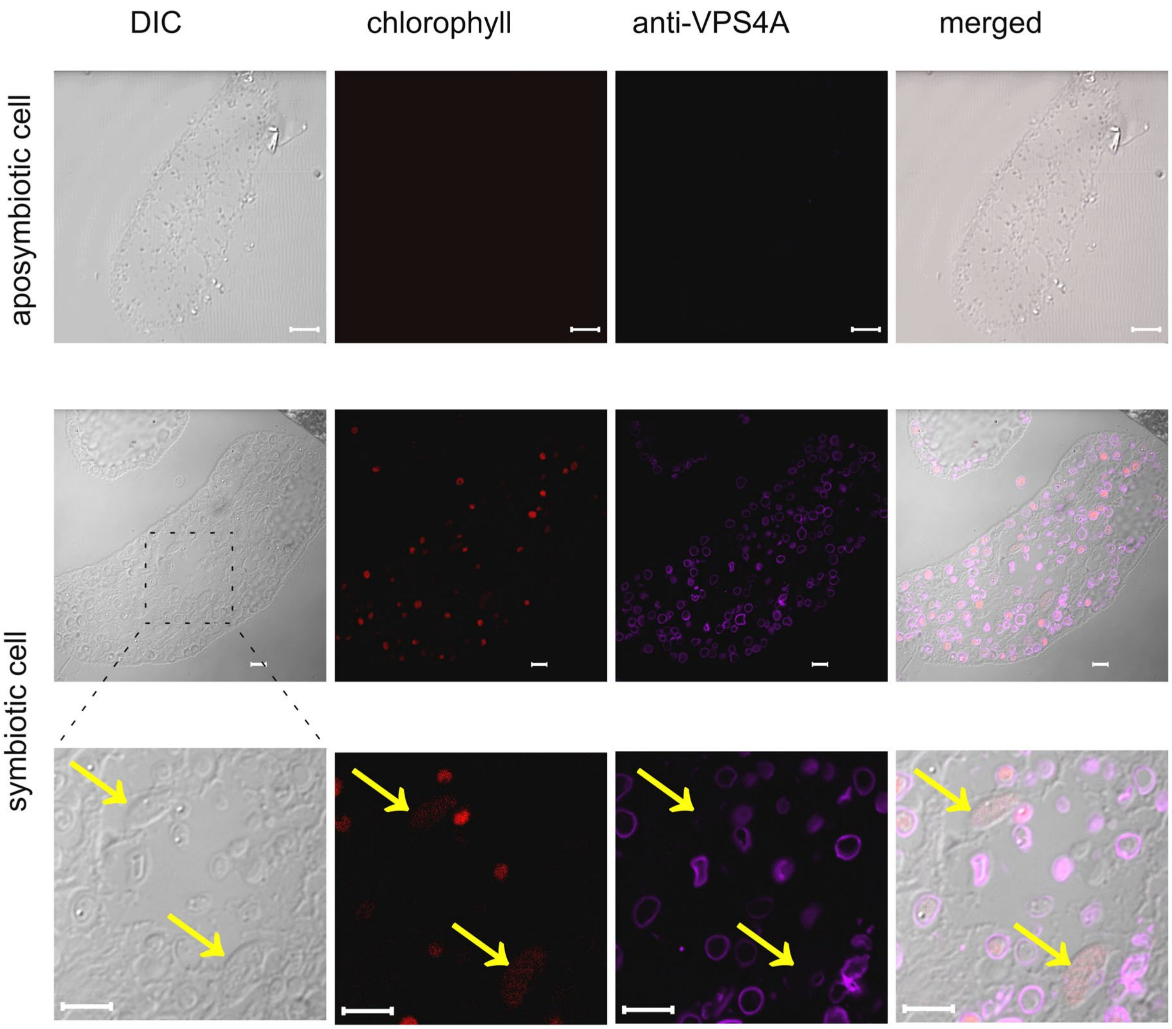


**Fig. S3. VPS4A localizes specifically to the perialgal vacuole membrane in symbiotic *P. bursaria* cells.**

Immunofluorescence images of aposymbiotic (top row) and symbiotic (middle and bottom rows) *P. bursaria* cells stained with anti-VPS4A antibody. Columns show DIC (differential interference contrast), chlorophyll autofluorescence (marking algal cells), anti-VPS4A immunofluorescence, and merged overlay. No VPS4A signal above background was detected in aposymbiotic cells, consistent with the dispersal of cluster 18 proteins across multiple subcellular compartments in aposymbiotic cells (Fig. 3A) and the absence of perialgal vacuoles. In symbiotic cells, VPS4A forms distinct ring-like structures surrounding endosymbiotic Chlorella cells (middle row). The boxed region is shown at higher magnification in the bottom row. Yellow arrows indicate ingested food algae (*Chlorogonium capillatum*) undergoing digestion in digestive vacuoles; no VPS4A ring is present around these cells, demonstrating that VPS4A ring formation is specific to the perialgal vacuole membrane and does not occur on digestive vacuole membranes. Scale bar, 5 μm

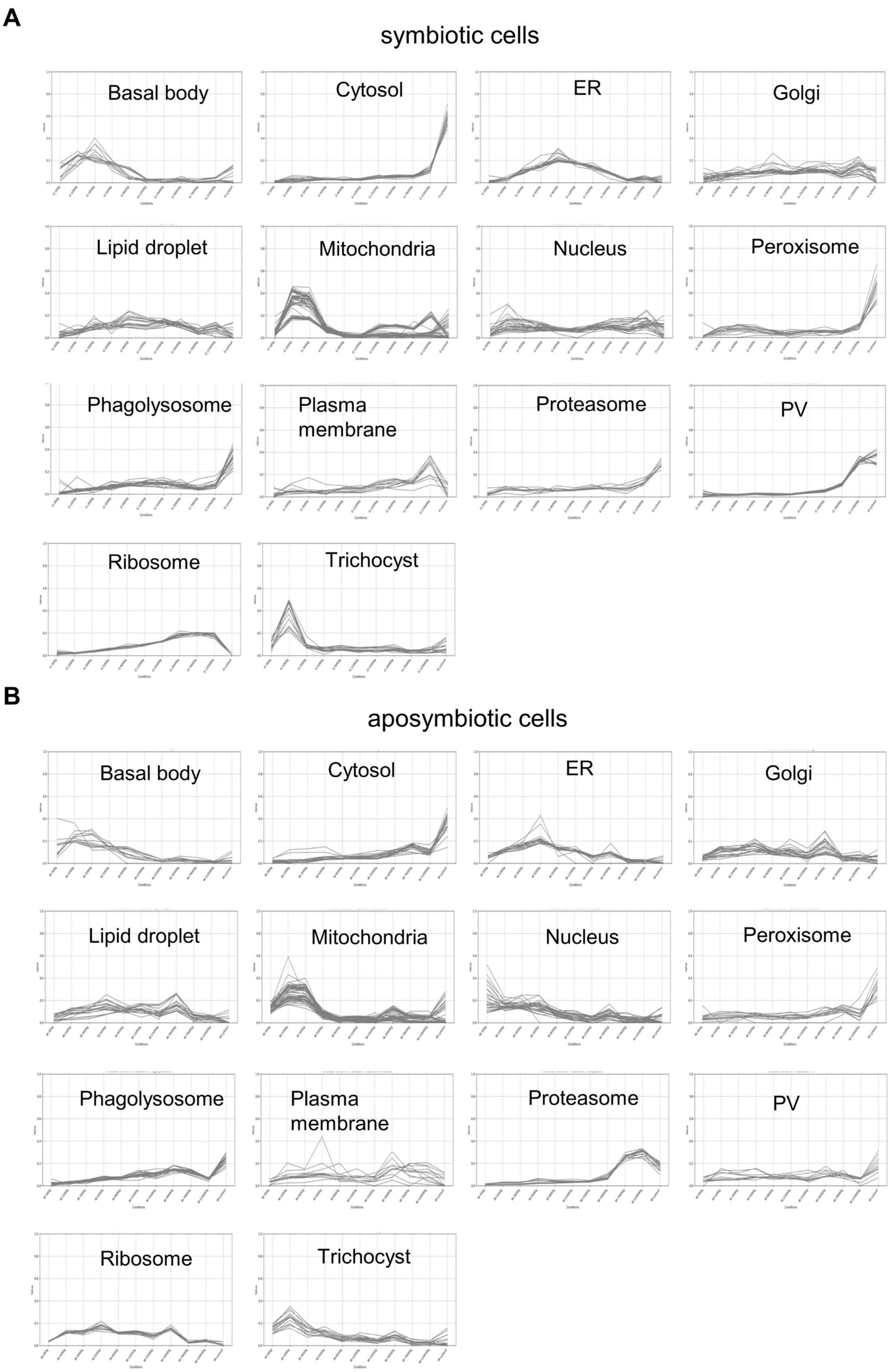
A
symbiotic cells
Basal body
Cytosol
ER
Golgi
Lipid droplet
Mitochondria
Nucleus
Peroxisome
Phagolysosome
Plasma membrane
Proteasome
PV
Ribosome
Trichocyst
B
aposymbiotic cells
Basal body
Cytosol
ER
Golgi
Lipid droplet
Mitochondria
Nucleus
Peroxisome
Phagolysosome
Plasma membrane
Proteasome
PV
Ribosome
Trichocyst

**Fig. S4. Fractionation distribution profiles of 252 subcellular compartment marker proteins.**

Profiles of each marker protein averaged from three replicates of symbiotic **(A)** and aposymbiotic **(B)** cells are shown.

**A**

**symbiotic cells**

Total Accuracy on Training Data: 0.9523809523809523
Classification Report for Each Marker:

| | precision | recall | f1-score | support |
|---|---|---|---|---|
| Basal body | 1.00 | 1.00 | 1.00 | 13 |
| Cytosol | 0.90 | 1.00 | 0.95 | 19 |
| ER | 0.81 | 1.00 | 0.89 | 17 |
| Golgi | 0.86 | 0.86 | 0.86 | 21 |
| Lipid droplet | 0.83 | 0.75 | 0.79 | 20 |
| Mitochondria | 1.00 | 1.00 | 1.00 | 54 |
| Nuclear | 0.96 | 1.00 | 0.98 | 27 |
| PV | 1.00 | 1.00 | 1.00 | 10 |
| Peroxisome | 1.00 | 0.85 | 0.92 | 13 |
| Phagolysosome | 1.00 | 1.00 | 1.00 | 22 |
| Plasma membrane | 1.00 | 0.75 | 0.86 | 12 |
| Proteasome | 1.00 | 1.00 | 1.00 | 16 |
| Ribosome | 1.00 | 1.00 | 1.00 | 14 |
| Trichocyst matrix | 1.00 | 1.00 | 1.00 | 15 |
| accuracy | | | 0.95 | 273 |
| macro avg | 0.95 | 0.94 | 0.95 | 273 |
| weighted avg | 0.95 | 0.95 | 0.95 | 273 |

**aposymbiotic cells**

Total Accuracy on Training Data: 0.9157509157509157
Classification Report for Each Marker:

| | precision | recall | f1-score | support |
|---|---|---|---|---|
| Basal body | 1.00 | 1.00 | 1.00 | 13 |
| Cytosol | 0.80 | 0.84 | 0.82 | 19 |
| ER | 0.77 | 1.00 | 0.87 | 17 |
| Golgi | 0.74 | 0.81 | 0.77 | 21 |
| Lipid droplet | 0.82 | 0.70 | 0.76 | 20 |
| Mitochondria | 1.00 | 1.00 | 1.00 | 54 |
| Nuclear | 1.00 | 1.00 | 1.00 | 27 |
| PV | 1.00 | 0.60 | 0.75 | 10 |
| Peroxisome | 0.82 | 0.69 | 0.75 | 13 |
| Phagolysosome | 0.88 | 1.00 | 0.94 | 22 |
| Plasma membrane | 1.00 | 0.83 | 0.91 | 12 |
| Proteasome | 1.00 | 1.00 | 1.00 | 16 |
| Ribosome | 1.00 | 1.00 | 1.00 | 14 |
| Trichocyst matrix | 1.00 | 1.00 | 1.00 | 15 |
| accuracy | | | 0.92 | 273 |
| macro avg | 0.92 | 0.89 | 0.90 | 273 |
| weighted avg | 0.92 | 0.92 | 0.91 | 273 |

**B**

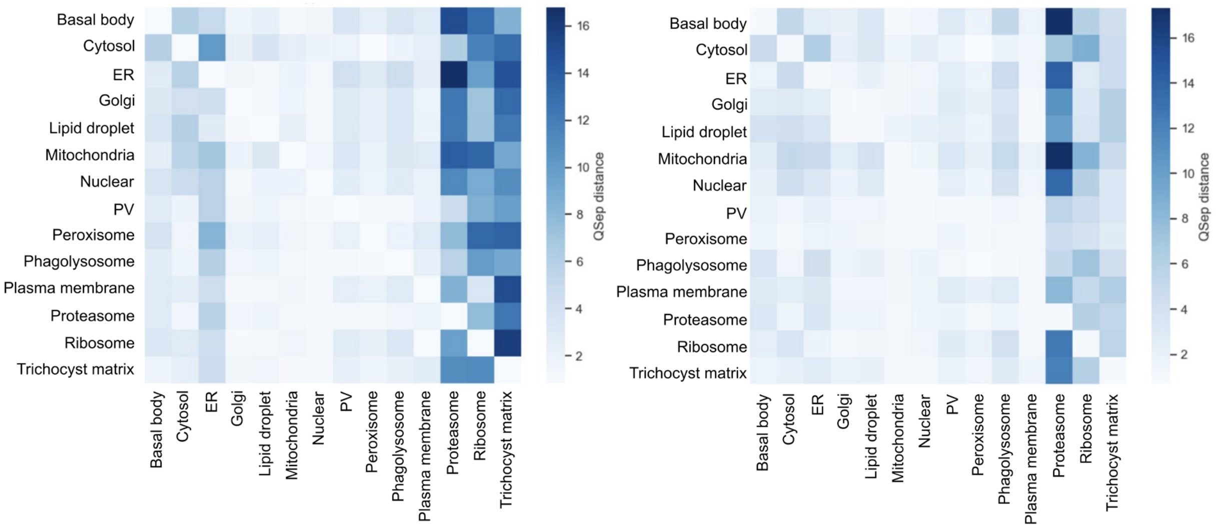


**Fig. S5. Quality examination of the marker proteins.**

**(A)** Prediction accuracy for marker proteins, as indicated by the F1 score for symbiotic and aposymbiotic cells. Organelle-specific recall indicates the proportion of markers correctly assigned to the cluster, and organelle-specific precision indicates the ratio of markers correctly assigned to the number of all markers assigned to the cluster. The F1 score is derived from the harmonic mean of recall and precision, representing the primary readout of SVM performance.

**(B)** Q-Sep analysis shows that proteins within the same subcellular compartment have shorter Q-Sep distances than those between different compartments.

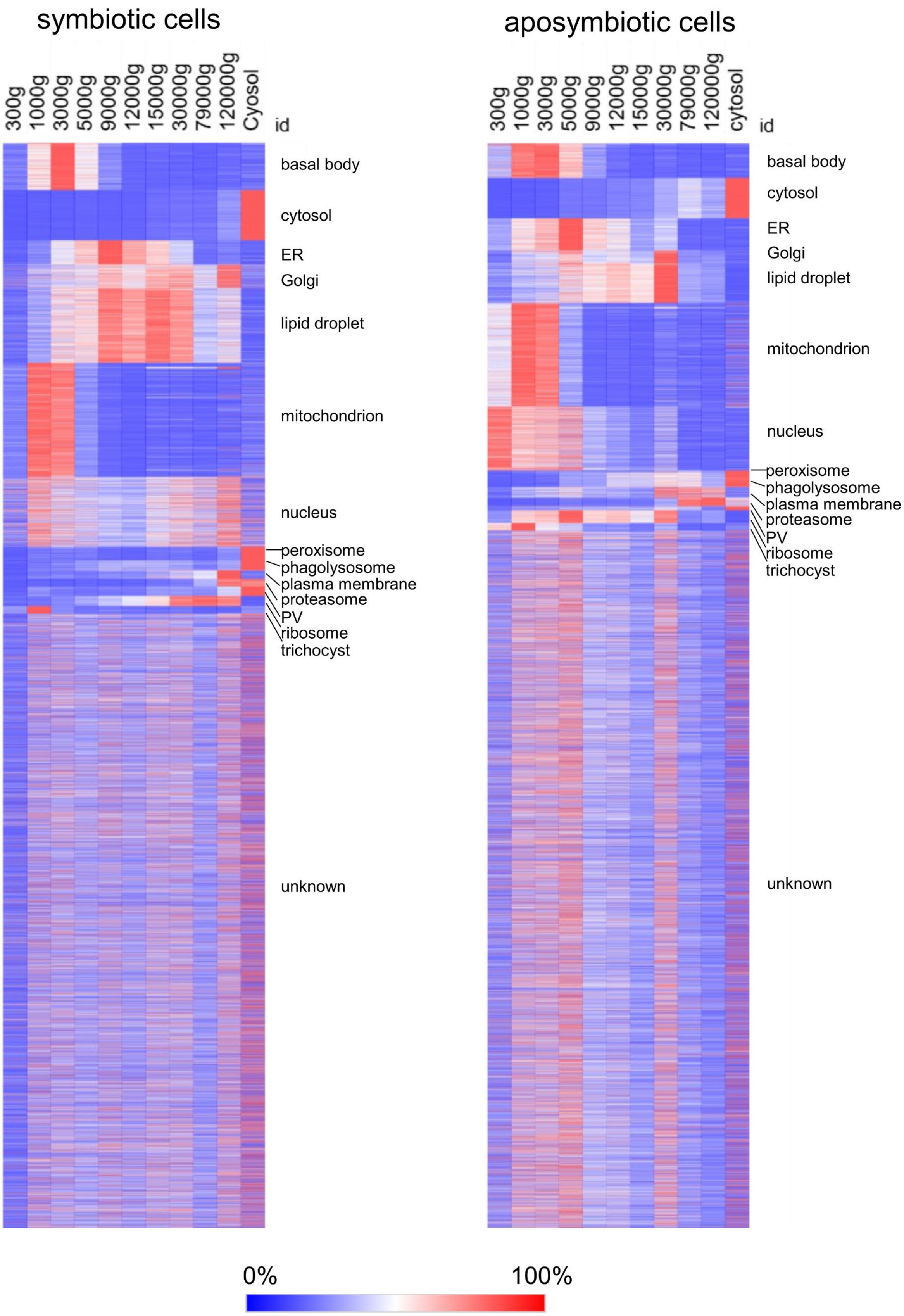
symbiotic cells
aposymbiotic cells
300g
1000g
3000g
5000g
9000g
12000g
15000g
30000g
79000g
120000g
Cyosol
cytosol
id
basal body
cytosol
ER
Golgi
lipid droplet
mitochondrion
nucleus
peroxisome
phagolysosome
plasma membrane
proteasome
PV
ribosome
trichocyst
unknown
0%
100%

**Fig. S6. Proteins from the same subcellular compartment show similar fractionation distribution profiles.**

Profiles of each protein averaged from three replicates are shown. The color intensity indicates the percentage of an individual protein contained in each fraction.

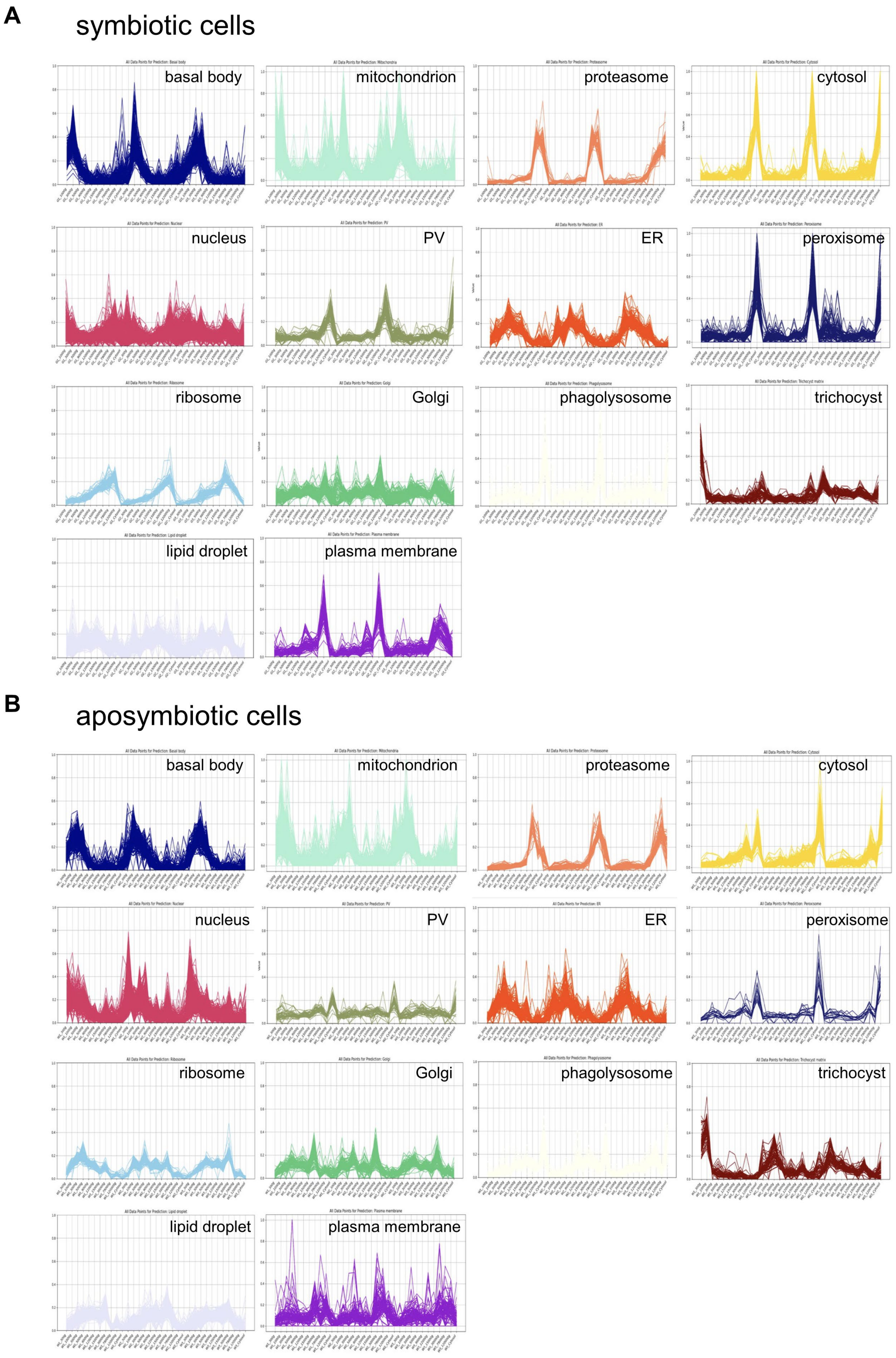
A
symbiotic cells
basal body
mitochondrion
proteasome
cytosol
nucleus
PV
ER
peroxisome
ribosome
Golgi
phagolysosome
trichocyst
lipid droplet
plasma membrane
B
aposymbiotic cells
basal body
mitochondrion
proteasome
cytosol
nucleus
PV
ER
peroxisome
ribosome
Golgi
phagolysosome
trichocyst
lipid droplet
plasma membrane

**Fig. S7. Fractionation distribution profiles of all proteins with predicted subcellular localizations.**

Concatenated profiles of each protein from three replicates of symbiotic **(A)** and aposymbiotic **(B)** cells are shown.

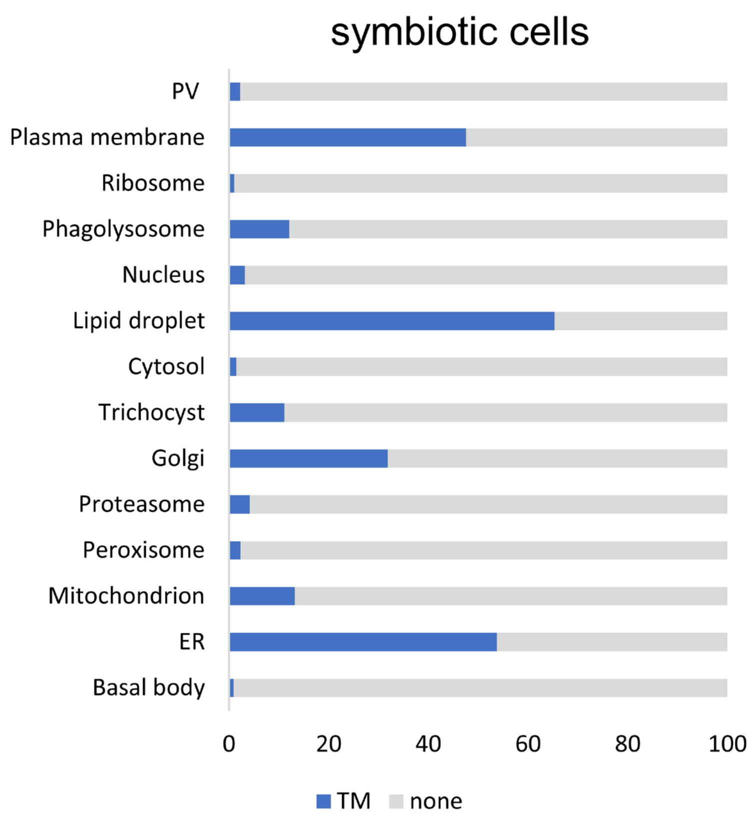
symbiotic cells
PV
Plasma membrane
Ribosome
Phagolysosome
Nucleus
Lipid droplet
Cytosol
Trichocyst
Golgi
Proteasome
Peroxisome
Mitochondrion
ER
Basal body
0
20
40
60
80
100
TM
none

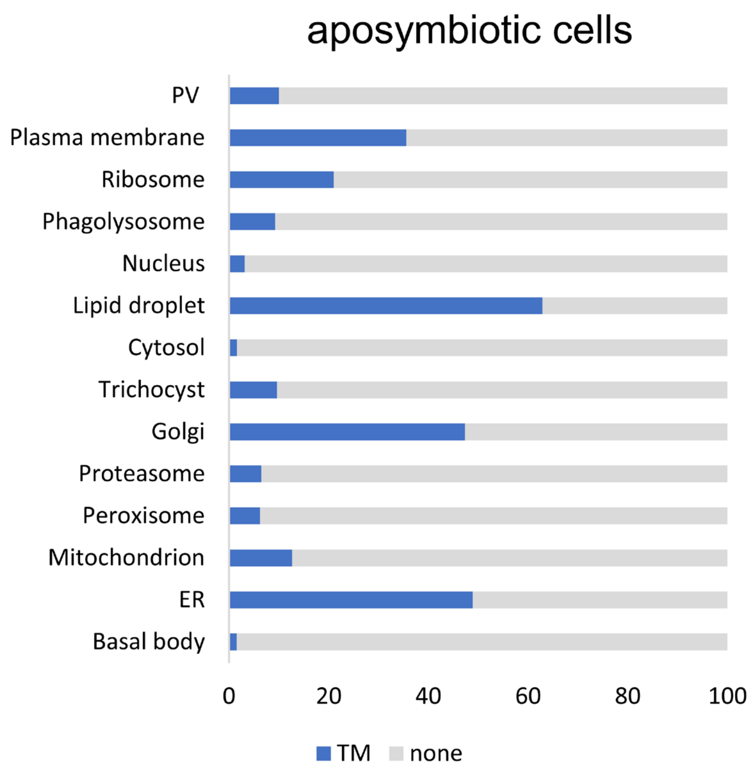
aposymbiotic cells
PV
Plasma membrane
Ribosome
Phagolysosome
Nucleus
Lipid droplet
Cytosol
Trichocyst
Golgi
Proteasome
Peroxisome
Mitochondrion
ER
Basal body
0
20
40
60
80
100
TM
none

**Fig. S8. Predicted potential transmembrane proteins using DeepTMHMM.**

The proportion of predicted transmembrane proteins in each subcellular compartment is shown for symbiotic (top) and aposymbiotic (bottom) *P. bursaria* cells.

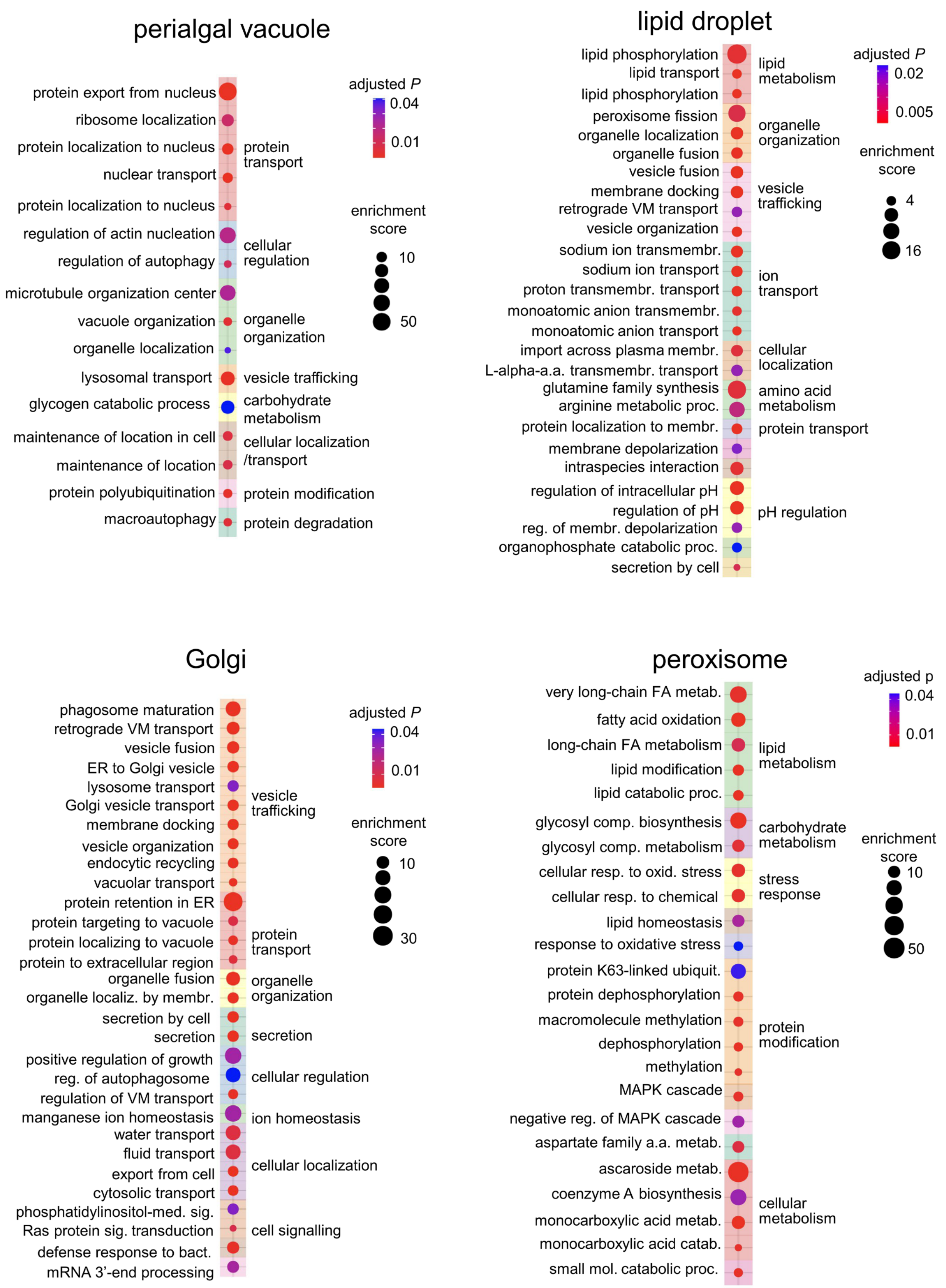

perialgal vacuole
protein export from nucleus
ribosome localization
protein localization to nucleus
nuclear transport
protein localization to nucleus
regulation of actin nucleation
regulation of autophagy
microtubule organization center
vacuole organization
organelle localization
lysosomal transport
glycogen catabolic process
maintenance of location in cell
maintenance of location
protein polyubiquitination
macroautophagy
protein transport
cellular regulation
organelle organization
vesicle trafficking
carbohydrate metabolism
cellular localization /transport
protein modification
protein degradation
adjusted P
0.04
0.01
enrichment score
10
50
lipid droplet
lipid phosphorylation
lipid transport
lipid phosphorylation
peroxisome fission
organelle localization
organelle fusion
vesicle fusion
membrane docking
retrograde VM transport
vesicle organization
sodium ion transmembr.
sodium ion transport
proton transmembr. transport
monoatomic anion transmembr.
monoatomic anion transport
import across plasma membr.
L-alpha-a.a. transmembr. transport
glutamine family synthesis
arginine metabolic proc.
protein localization to membr.
membrane depolarization
intraspecies interaction
regulation of intracellular pH
regulation of pH
reg. of membr. depolarization
organophosphate catabolic proc.
secretion by cell
lipid metabolism
organelle organization
vesicle trafficking
ion transport
cellular localization
amino acid metabolism
protein transport
pH regulation
adjusted P
0.02
0.005
enrichment score
4
16
Golgi
phagosome maturation
retrograde VM transport
vesicle fusion
ER to Golgi vesicle
lysosome transport
Golgi vesicle transport
membrane docking
vesicle organization
endocytic recycling
vacuolar transport
protein retention in ER
protein targeting to vacuole
protein localizing to vacuole
protein to extracellular region
organelle fusion
organelle localiz. by membr.
secretion by cell
secretion
positive regulation of growth
reg. of autophagosome
regulation of VM transport
manganese ion homeostasis
water transport
fluid transport
export from cell
cytosolic transport
phosphatidylinositol-med. sig.
Ras protein sig. transduction
defense response to bact.
mRNA 3'-end processing
vesicle trafficking
protein transport
organelle organization
secretion
cellular regulation
ion homeostasis
cellular localization
cell signalling
adjusted P
0.04
0.01
enrichment score
10
30
peroxisome
very long-chain FA metab.
fatty acid oxidation
long-chain FA metabolism
lipid modification
lipid catabolic proc.
glycosyl comp. biosynthesis
glycosyl comp. metabolism
cellular resp. to oxid. stress
cellular resp. to chemical
lipid homeostasis
response to oxidative stress
protein K63-linked ubiquit.
protein dephosphorylation
macromolecule methylation
dephosphorylation
methylation
MAPK cascade
negative reg. of MAPK cascade
aspartate family a.a. metab.
ascaroside metab.
coenzyme A biosynthesis
monocarboxylic acid metab.
monocarboxylic acid catab.
small mol. catabolic proc.
lipid metabolism
carbohydrate metabolism
stress response
protein modification
cellular metabolism
adjusted p
0.04
0.01
enrichment score
10
50

**Fig. S9. Gene ontology (GO) analysis of symbiotic cell-specific compartment proteins.** Gene Ontology Biological Process (GOBP) analysis of symbiotic cell-specific proteins in the PV, lipid droplet, Golgi, and peroxisome. These organelles showed significant changes in protein numbers between symbiotic and aposymbiotic cells. Only terms with an enrichment score greater than 2 and an adjusted P value less than 0.05 are selected (see also data file S7A). Non-redundant specific terms are listed on the left and general parent terms are listed on the right.

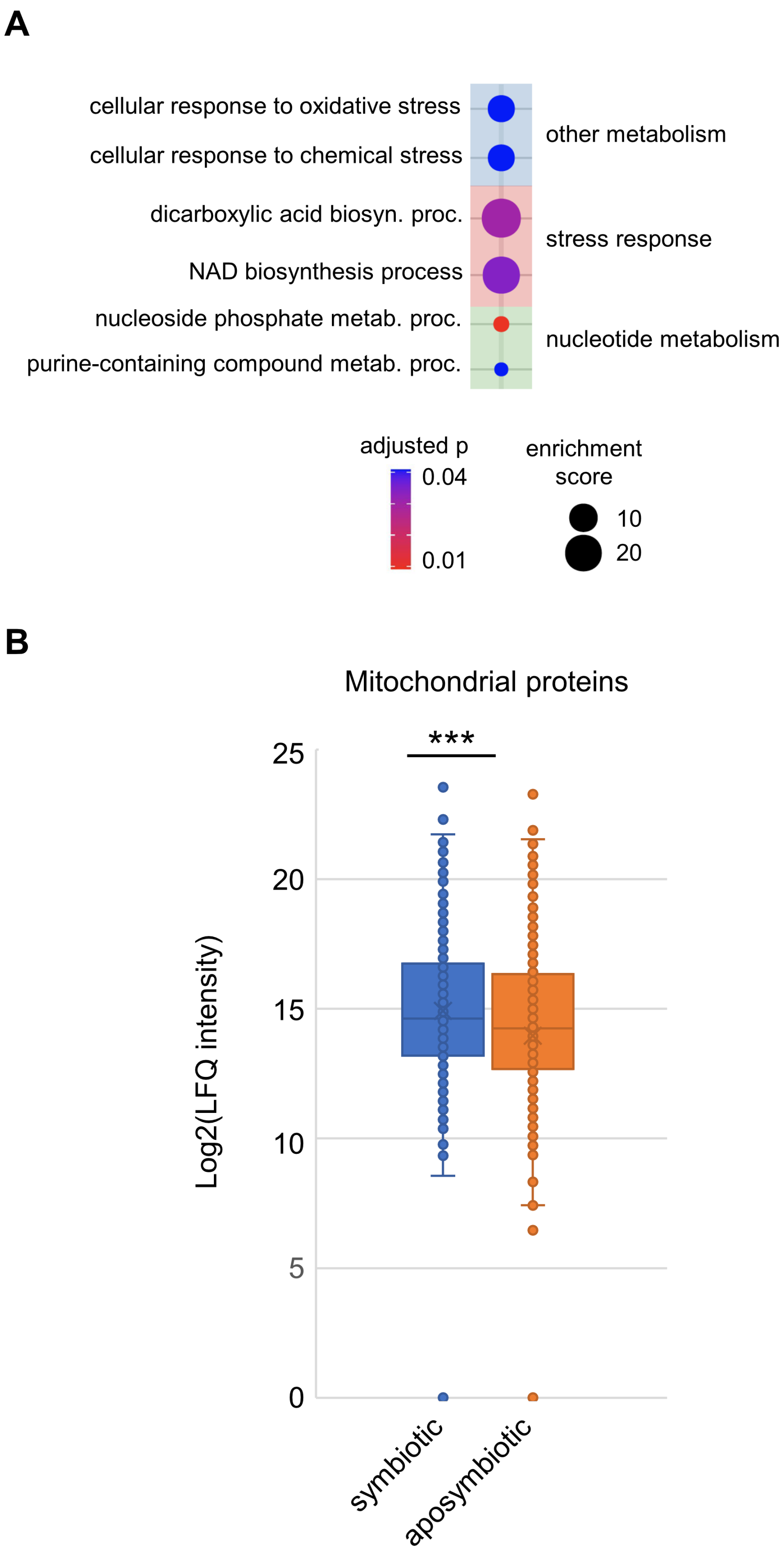
A
cellular response to oxidative stress
cellular response to chemical stress
dicarboxylic acid biosyn. proc.
NAD biosynthesis process
nucleoside phosphate metab. proc.
purine-containing compound metab. proc.
other metabolism
stress response
nucleotide metabolism
adjusted p
0.04
0.01
enrichment score
10
20
B
Mitochondrial proteins
***
Log2(LFQ intensity)
25
20
15
10
5
0
symbiotic
aposymbiotic

**Fig. S10. GO analysis of symbiotic cell-specific mitochondrial proteins and the abundance of mitochondrial proteins in symbiotic and aposymbiotic cells.**

**(A)** GOBP analysis of symbiotic cell-specific proteins in mitochondria. Only terms with an enrichment score greater than 2 and an adjusted P value less than 0.05 are selected (see also data file S7A). Non-redundant specific terms are listed on the left and general parent terms are listed on the right. **(B)** Mitochondrial protein abundance in symbiotic cells is higher than that in aposymbiotic cells. Protein abundance data were obtained from total lysate proteomes (data file S2) and the abundance of 780 proteins that were detected in both symbiotic and aposymbiotic cells was compared. ***, $P$ value < 0.001, Paired-sample t-test, n = 780 proteins.

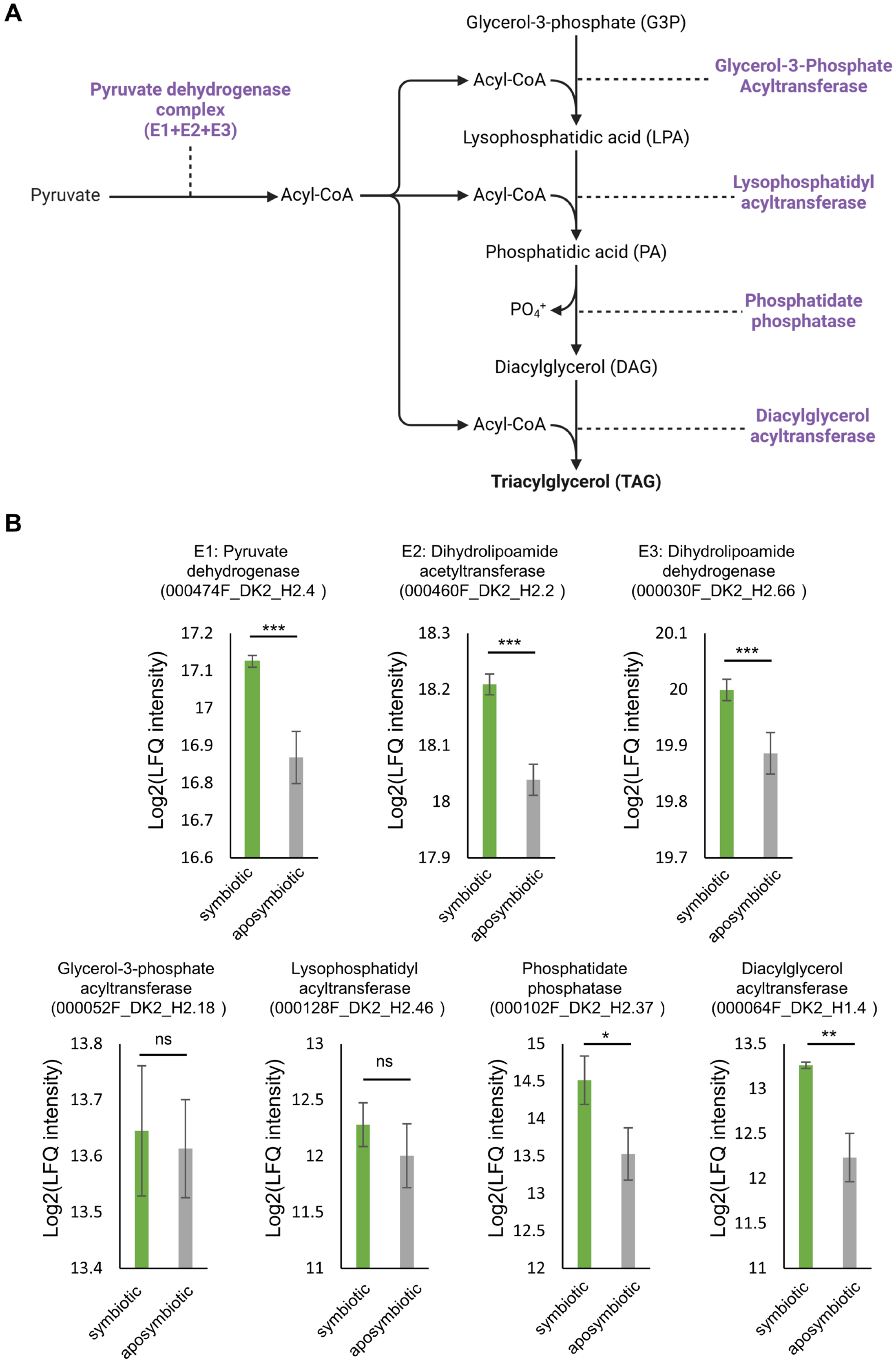
A
Glycerol-3-phosphate (G3P)
Pyruvate dehydrogenase complex (E1+E2+E3)
Pyruvate
Acyl-CoA
Acyl-CoA
Glycerol-3-Phosphate Acyltransferase
Lysophosphatidic acid (LPA)
Acyl-CoA
Lysophosphatidyl acyltransferase
Phosphatidic acid (PA)
PO4+
Phosphatidate phosphatase
Diacylglycerol (DAG)
Acyl-CoA
Diacylglycerol acyltransferase
Triacylglycerol (TAG)
B
E1: Pyruvate dehydrogenase (000474F_DK2_H2.4 )
E2: Dihydrolipoamide acetyltransferase (000460F_DK2_H2.2 )
E3: Dihydrolipoamide dehydrogenase (000030F_DK2_H2.66 )
Glycerol-3-phosphate acyltransferase (000052F_DK2_H2.18 )
Lysophosphatidyl acyltransferase (000128F_DK2_H2.46 )
Phosphatidate phosphatase (000102F_DK2_H2.37 )
Diacylglycerol acyltransferase (000064F_DK2_H1.4 )
Log2(LFQ intensity)
symbiotic
aposymbiotic
***
ns
*
**

**Fig. S11. Several enzymes involved in triacylglycerol (TAG) synthesis increase their protein abundance in symbiotic *P. bursaria* cells.**

**(A)** The biosynthesis pathway of triacylglycerol. **(B)** Enzymes involved in triacylglycerol synthesis are increased in symbiotic cells. Protein abundance data were obtained from total lysate proteomes (data file S2). *, $P$ value < 0.05; **, $P$ value < 0.01; ***, $P$ value < 0.001; N.S., not significant; Student's t-test, n = 3.

**Data files**

**Data file S1. Fractionation mass spectrometry data for symbiotic and aposymbiotic *Paramecium bursaria* cells**. *P. bursaria* cells were lysed by nitrogen cavitation and separated into 11 fractions by differential centrifugation (300 × g, 1,000 × g, 3,000 × g, 5,000 × g, 9,000 × g, 12,000 × g, 15,000 × g, 30,000 × g, 79,000 × g, 120,000 × g, and cytosolic supernatant). Three biological replicates of symbiotic (G1-G3) and aposymbiotic (W1-W3) cells were analyzed by DIA-LC/MS using a timsTOF HT mass spectrometer, and data were processed with Spectronaut. Proteins not detected in all three replicates of either cell type were excluded. **(A)** Raw MS intensity values for 11,182 protein groups across all 66 fractionation samples (6 cell-replicate combinations × 11 fractions). Missing values indicate proteins not detected in that fraction. **(B)** Sum-normalized abundance profiles (values scaled to sum to 1 per protein) used as input for HDBSCAN unsupervised clustering and SVM-based subcellular localization prediction.

**Data file S2. Proteomic data from total cell lysates.** Label-free quantification (LFQ) intensities for 6,586 protein groups quantified across three biological replicates of symbiotic (DK2g-1, DK2g-2, DK2g-3) and aposymbiotic (DK2w-1, DK2w-2, DK2w-3) *P. bursaria* cells from total cell lysate samples. Protein abundance differences between symbiotic and aposymbiotic cells were evaluated by Student's t-test with Benjamini-Hochberg correction ($P < 0.05$).

**Data file S3.** H**DBSCAN unsupervised clustering and GO cellular component (GOCC) enrichment results.** Normalized protein fractionation profiles from 33 fractions (11 fractions × 3 replicates) were analyzed by hierarchical density-based spatial clustering of applications with noise (HDBSCAN) implemented in Python Scikit-learn. The algorithm generated 20 clusters in symbiotic cells and 19 clusters in aposymbiotic cells; proteins with ambiguous profiles are labeled "unclustered." **(A)** Cluster assignments for 10,255 proteins detected in symbiotic *P. bursaria* cells (green_hdbscan_cluster column). **(B)** Cluster assignments for 10,255 proteins detected in aposymbiotic *P. bursaria* cells (hdbscan_cluster column). **(C)** GOCC enrichment analysis (clusterProfiler, BH-adjusted $P < 0.05$) of each HDBSCAN cluster in symbiotic cells, including cluster number, cluster label, GO term ID, GO term description, gene ratio, background ratio, fold enrichment, p-value, adjusted p-value, q-value, gene count, and constituent gene IDs. **(D)** GOCC enrichment analysis of each HDBSCAN cluster in aposymbiotic cells.

**Data file S4. Marker proteins and SVM-based subcellular localization predictions**. A total of 252 marker proteins spanning 14 subcellular compartments were curated based on ortholog information from *Paramecium tetraurelia*, InterProScan annotations, and HDBSCAN clustering. The 14 compartments are: cytosol, proteasome, ribosome, peroxisome, endoplasmic reticulum (ER), Golgi, plasma membrane, nucleus, lipid droplet, basal body, mitochondrion, phagolysosome, perialgal vacuole (PV), and trichocyst. SVM was trained using these markers (F1 scores: 95.2% for symbiotic, 91.5% for aposymbiotic cells). **(A)** Curated marker protein list with compartment assignment, identification method, and literature reference. **(B)** SVM-predicted compartment assignment probabilities for 9,982 proteins quantified in symbiotic *P. bursaria* cells, including probabilities for each of the 14 compartments, confidence level (very high: >0.9, high: >0.8, medium: >0.5), maximum probability value, and the most probable compartment class. **(C)** SVM-predicted compartment assignment probabilities for 9,982 proteins in aposymbiotic *P. bursaria* cells. **(D)** GOCC enrichment analysis (clusterProfiler, BH-adjusted $P < 0.05$) of all curated marker protein sets across the 14 compartments. **(E)** GOCC enrichment analysis of SVM-classified proteins (medium confidence or higher) in symbiotic *P. bursaria* cells, grouped by predicted compartment. **(F)** GOCC enrichment analysis of SVM-classified proteins in aposymbiotic *P. bursaria* cells.

**Data file S5. t-SNE dimensionality reduction of fractionation proteomics data**. Two-dimensional t-SNE (t-distributed stochastic neighbor embedding) coordinates were computed from the 33-dimensional fractionation profiles (11 fractions × 3 replicates per cell type) for 10,255 proteins detected in both cell types, using perplexity = 60 and a maximum of 1,000 iterations. **(A)** t-SNE coordinates for proteins from aposymbiotic *P. bursaria* cells. **(B)** t-SNE coordinates for proteins from symbiotic *P. bursaria* cells.

**Data file S6. Deep learning-based predictions of protein subcellular localization and transmembrane topology for all 10,255 proteins detected in *P. bursaria*.** **(A)** DeepLoc 2.0 subcellular localization prediction scores across 10 compartment categories (cell membrane, cytoplasm, endoplasmic reticulum, extracellular, Golgi apparatus, lysosome/vacuole, mitochondrion, nucleus, peroxisome, and plastid), with the final predicted localization for each protein. **(B)** DeepTMHMM transmembrane topology predictions, including the predicted number of transmembrane helices (Predictions column), topology classification (TM: transmembrane protein; nTM: non-transmembrane protein), and label name.

**Data file S7. Enrichment analysis of symbiotic cell-specific organelle proteins**. Proteins uniquely assigned to the perialgal vacuole (PV), lipid droplet, Golgi apparatus, peroxisome, and mitochondria in symbiotic cells (i.e., absent or substantially reduced in aposymbiotic cells) were subjected to enrichment analysis. **(A)** Gene Ontology Biological Process (GOBP) enrichment results, including GO term ID, gene IDs, GO term description, biological categories, user-defined categories, BH-adjusted p-value, q-value, gene count, and enrichment score. Only terms with enrichment score > 2 and adjusted $P < 0.05$ are included. **(B)** KEGG pathway enrichment results for symbiotic cell-specific proteins in the same organelles.

**Data file S8.** Q-Sep distances between all pairs of subcellular compartments. Q-Sep quantifies the quality of spatial proteomics separation by calculating the ratio of inter-compartment to intra-compartment distance distributions for SVM-predicted protein sets; larger Q-Sep values indicate cleaner separation between compartments. Values range from 0 (complete overlap) to 1 (perfect separation). **(A)** Q-Sep distance matrix for 13 subcellular compartments in symbiotic *P. bursaria* cells. **(B)** Q-Sep distance matrix for 13 subcellular compartments in aposymbiotic *P. bursaria* cells.
